# A single epigenetic repressor controls opposing developmental tempos in human neural and cranial neural crest lineages

**DOI:** 10.64898/2026.10.09.758088

**Authors:** Zeynep Eda Atmaca Duran, Marcella Birtele, Shweta Mendiratta, Ramesh Yelgandula, Claire Stevens, Suhn Rhie, Giorgia Quadrato, Oliver Bell

**Affiliations:** Department of Cancer Biology, Keck School of Medicine, University of Southern California, Los Angeles, CA, 90033, USA; Department of Stem Cell Biology and Regenerative Medicine, Keck School of Medicine, University of Southern California, Los Angeles, CA, 90033, USA; Eli and Edythe Broad CIRM Center for Regenerative Medicine and Stem Cell Research at USC, Keck School of Medicine, University of Southern California, Los Angeles, CA, 90033, USA; Laboratory of Epigenetics, Cell Fate and Disease, Centre for DNA Fingerprinting and Diagnostics, Hyderabad, India

## Abstract

The precise timing of lineage specification is essential for embryonic development, yet the epigenetic mechanisms enforcing human developmental tempo remain poorly understood^1,2^. Heterozygous loss-of-function variants in *ZNF462* cause Weiss-Kruszka syndrome (WKS), a pleiotropic disorder characterized by neurodevelopmental delays and craniofacial dysmorphisms^3–5^. Here, we show that ZNF462 acts as an epigenetic gatekeeper that coordinates human developmental tempo across divergent lineages by repressing cell lineage-specific gene enhancers. In human embryonic stem cells (hESCs), ZNF462 maintains self-renewal by recruiting the G9A/GLP histone methyltransferase complex to silence primate-specific retrotransposon-derived enhancers. Upon neural lineage commitment, ZNF462 redirects to hominoid-specific LTR5_Hs elements and divergent lineage-specific enhancer programs, where its loss produces opposing kinetics. In neural progenitor cells (NPCs), ZNF462 enforces G9A/GLP-dependent repression of ZIC-driven neural enhancers, and loss of this repression traps NPCs in a progenitor state, slowing cortical neurogenesis without impairing terminal neurogenic capacity. In neural crest cells (NCCs), ZNF462 targets repressive chromatin to silence SOX9 family-driven enhancers. Here, ZNF462 haploinsufficiency triggers premature activation of definitive NCC specifiers, accelerating differentiation. These findings establish that a single repressor can simultaneously decelerate and accelerate developmental tempo by gating lineage-specific enhancers, placing Weiss-Kruszka syndrome into the emerging class of human developmental timing disorders.

## Introduction

During embryonic development, the precise timing of cell fate specification and tissue maturation is essential for the formation of functional organs^1,2^. Distinct cell lineages must execute their differentiation programs in a coordinated sequence, with transcriptional repressors playing a critical role in preventing premature activation of lineage-specific gene programs^6,7^. When this temporal coordination is disrupted, even by heterozygous mutations in a single gene, the consequences can extend across multiple organ systems^1,2,8,9^. Weiss-Kruszka syndrome (WKS; OMIM #618619) is a neurodevelopmental disorder characterized by global developmental delay, autism spectrum disorder (ASD), hypotonia, and distinct craniofacial abnormalities, including ptosis, down-slanting palpebral fissures, and metopic ridging^3–5^. The genetic etiology of this syndrome is driven by heterozygous loss-of-function mutations in the *ZNF462* gene. *ZNF462* encodes a ∼250 kDa, highly conserved, vertebrate-specific protein characterized by 34 C_2_H_2_-type zinc finger domains, yet its role in human neural development is unknown.

The pleiotropic features of WKS indicate a crucial function in coordinating two distinct lineages that diverge at the embryonic neural plate border: central nervous system precursors and cranial neural crest cells (NCCs). Lineage bifurcation at the neural plate border is orchestrated by multi-tiered transcription factor (TF) networks operating downstream of WNT, BMP, and FGF signaling^10–12^. Within this regulatory hierarchy, ZIC and SOX family factors represent major lineage-defining nodes^13–15^. ZIC2 and ZIC3 regulate neural progenitor potency and differentiation timing, whereas SOX9 and SOX10, acting alongside specifiers like TFAP2, TWIST, and SNAIL, govern neural crest specification and epithelial-to-mesenchymal transition (EMT)^16,17^.

Previous work demonstrated that the murine homolog *Zfp462* is essential for embryonic development and mouse ESC self-renewal, encoding a TF that safeguards neural lineage specification by recruiting the G9A/GLP methyltransferase complex to silence TE-derived enhancers^18–20^. However, because *Zfp462* targets are predominantly enriched at mouse-specific LTRs, the gene regulatory networks governed by ZNF462 in human cells are likely distinct^18^. Consequently, how ZNF462 engages human-specific regulatory elements to coordinate neural and neural crest-specific gene expression programs and how its haploinsufficiency disrupts the development of each lineage remains unknown.

Here, we engineered human ESCs (hESCs) recapitulating clinically relevant heterozygous *ZNF462* loss-of-function mutations and utilized directed NCC and neural progenitor cell (NPC) differentiation to model WKS pathogenesis. We demonstrate that ZNF462 is an epigenetic gatekeeper of human developmental tempo. ZNF462 enforces developmental timing by repressing lineage-specific enhancer programs. In NPCs, ZNF462 represses *ZIC2* and downstream neural crest border programs via G9A/GLP recruitment, and its loss traps NPCs in a progenitor state and slows cortical neurogenesis, establishing a developmental delay without compromising the intrinsic differentiation potential. In NCCs, ZNF462 mediates silencing of SOX9-driven enhancers, and its loss accelerates cranial neural crest specification and EMT. Together, our findings establish ZNF462 as an epigenetic gatekeeper that dictates human developmental tempo and uncover that haploinsufficiency alters the pace of lineage specification and maturation without losing cell fate competence, providing a unifying molecular basis for the pleiotropic defects in WKS.

## Results

### ZNF462 is required for hESC self-renewal by repressing neuroectodermal genes

To elucidate the mechanistic basis of WKS and determine the impact of *ZNF462* heterozygous loss-of-function mutations on human neural development, we used CRISPR-Cas9-assisted homology-directed repair to engineer clinically relevant truncating mutations into hESCs. Because WKS is driven by *de novo* mutations originating in germ cells or early embryonic development, modeling these variants in hESCs and downstream differentiated neural cell types where ZNF462 is highly expressed enabled us to capture the primary consequences of haploinsufficiency in disease-relevant developmental contexts (Ext. Data Fig. 1a). Pathogenic ZNF462 variants are predominantly truncating mutations within exon 3, which comprises 54% of the coding region (Fig. 1a). To capture the primary WKS clinical spectrum, we modeled two distinct truncating mutations. The p.Arg1263Ter (R1263*) mutation, previously identified in multiple members of a single family, is associated with variable craniofacial dysmorphisms, including metopic ridging, ptosis, arched eyebrows, and a small upturned nose, as well as severe structural brain abnormalities such as agenesis of the corpus callosum and colpocephaly (ClinVar: VCV000402116)^3,4^. The p.Tyr1200Ter (Y1200*) mutation was designed to model several frameshift variants leading to early termination, including p.Met1096ThrfsTer10 (M1096fs*10; ClinVar: VCV003341478.1), which is associated with autism spectrum disorder and speech delay, and p.Arg1168fsTer15 (R1168fs*15; ClinVar: VCV002500924.1), which is associated with craniofacial dysmorphisms and global developmental delay. We confirmed successful gene editing by Sanger sequencing and validated heterozygous loss-of-function by western blot analysis, which showed an approximately 50% reduction in full-length ZNF462 protein in both nonsense mutant lines (Fig. 1b, Ext. Data Fig. 1b and c).

**Fig. 1.**
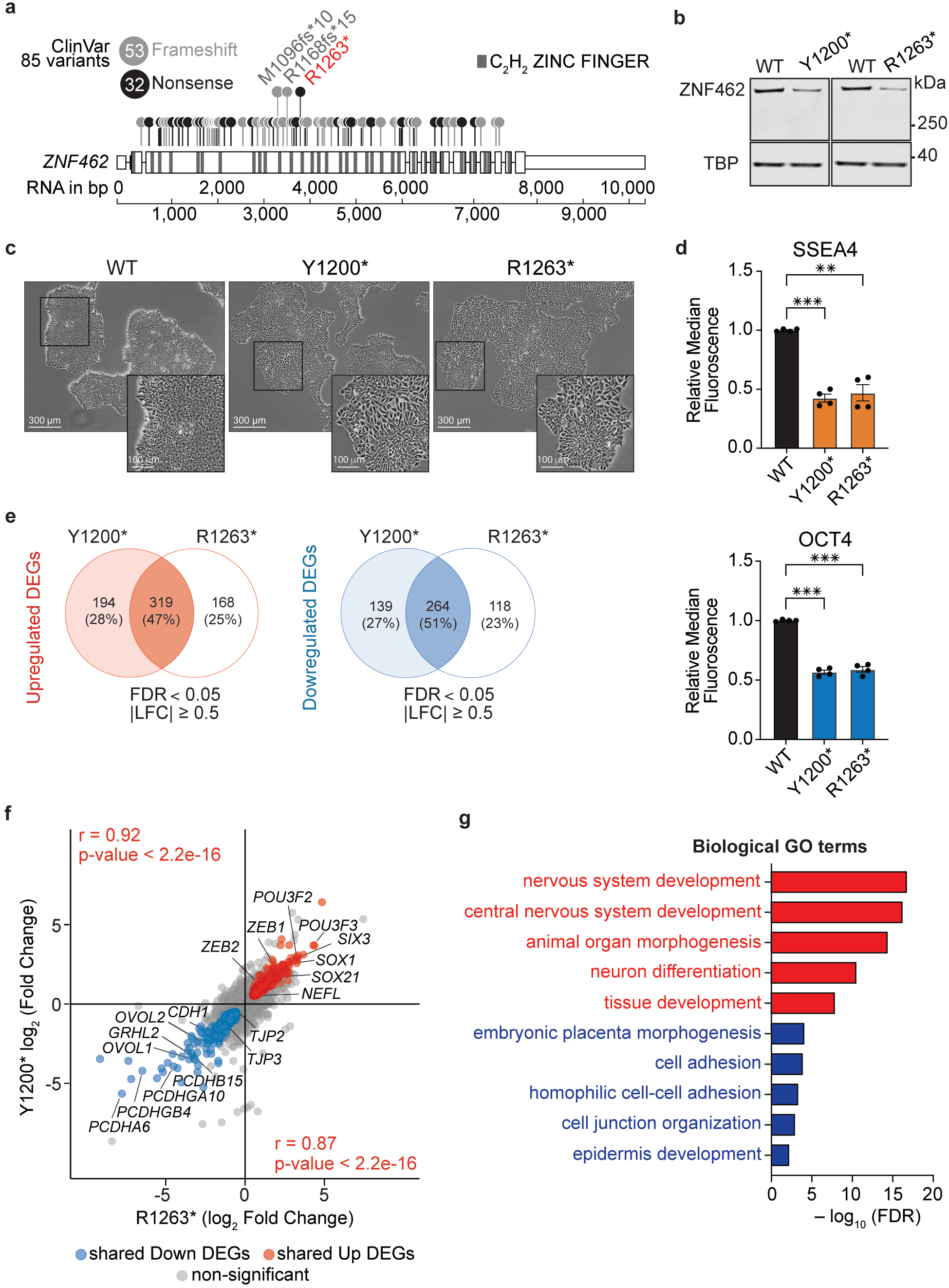
ZNF462 is required for silencing of neuroectodermal genes in hESC. **a)** Schematic representation of the *ZNF462* RNA transcript model with 85 ClinVar-reported truncating gene variants (frameshift and nonsense) mapped to exons (white boxes) and introns (black line). Domains encoding C_2_H_2_ zinc fingers are marked in dark grey. Highlighted are the pathogenic R1263* variant (red) and two ClinVar truncating variants located near the engineered Y1200* mutation (light grey). **b)** Western blot of ZNF462 protein levels in wild-type (WT) and two independent *ZNF462* mutant hESC lines, Y1200* and R1263*. TBP serves as a loading control. **c)** Representative phase-contrast images of WT, Y1200*, and R1263* hESC colonies. Insets show higher-magnification views of colony morphology. **d)** Flow cytometry quantification of SSEA4 and OCT3/4 expression in Y1200* and R1263* hESCs relative to WT. Bars represent mean ± SEM from n = 4 independent matched experiments. Statistical significance was assessed by one-way repeated-measures ANOVA with Greenhouse-Geisser correction followed by Dunnett’s multiple comparisons test using WT as the reference group. **p < 0.01, ***p < 0.001. **e)** Venn diagrams of significantly up- and downregulated DEGs in Y1200* and R1263* hESCs compared with WT (FDR < 0.05 and |log_2_ fold change| ≥ 0.5). **f)** Pairwise comparison of R1263* and Y1200* DEGs. Shared upregulated genes are highlighted in red, and shared downregulated genes in blue. Pearson correlation coefficients and corresponding P values are shown. **g)** Gene Ontology (GO) analysis of shared Y1200* and R1263* DEGs. Bars show significantly enriched biological-process GO terms for shared upregulated (red) and downregulated (blue) DEGs; significance is shown as -log_10_ (FDR).

Both mutant hESC lines displayed substantial morphological differences compared with wild-type hESCs, including large, flattened cells spreading outward from dense hESC colonies (Fig. 1c). Flow cytometry further demonstrated significantly reduced expression of the pluripotency markers SSEA4 and OCT4, consistent with spontaneous differentiation and suggesting that ZNF462 is required for the maintenance of hESC self-renewal (Fig. 1d, Ext. Data Fig. 1d). To investigate the gene expression changes underlying this self-renewal defect, we performed bulk RNA-seq which identified 916 and 869 significant differentially expressed genes (DEGs; false discovery rate (FDR) < 0.05; |L2FC| ≥ 0.5) in the Y1200* and R1263* lines, respectively (Ext. Data Fig. 1e, f; Ext. Data Table 1). DEGs from both mutant lines were highly correlated and largely overlapping, indicating a shared transcriptional consequence of ZNF462 truncation in hESCs (Fig. 1e, f). Shared DEGs indicated a coordinated shift in cellular identity with upregulation of early neuroectoderm regulators (*SOX1, SOX21, SIX3, POU3F2/BRN2, NEFL*) and EMT markers (*ZEB1/2*), and downregulation of epithelial and adhesion genes (*OVOL1/2, GRHL2, CDH1*). GO term analysis confirmed enrichment for nervous system development among upregulated DEGs and cell adhesion/epidermis development among downregulated DEGs (Fig. 1g). These findings indicate that ZNF462 maintains hESC identity by repressing early neuroectodermal differentiation programs, and its haploinsufficiency promotes premature neural differentiation.

### Heterozygous ZNF462 loss impairs targeting of G9A/GLP-dependent heterochromatin

To investigate how ZNF462 represses early neuroectodermal gene expression programs, we performed Cleavage Under Targets and Release Using Nuclease (CUT&RUN) profiling to identify direct ZNF462 targets and determine their overlap with G9A/GLP and H3K9me2 in wild-type hESCs. CUT&RUN ZNF462 profiling identified 7,844 high-confidence binding sites (Fig. 2a, Ext. Data Fig. 2a, Ext. Data Table 2). G9A and WIZ, the core subunit that scaffolds the histone methyltransferase complex, showed strong enrichment at ZNF462 targets, suggesting that the mechanism of repression is conserved between human and mouse ESCs (Fig. 2b, Ext. Data Fig. 2b)^18^. H3K9me2 showed more graded enrichment, suggesting context-dependent deposition or turnover. Using CUT&RUN with *Drosophila* spike-in normalization, we quantified changes in ZNF462 binding and heterochromatin in mutant hESCs. Both mutant lines showed significant reduction in ZNF462 enrichment at high-confidence binding sites including at *SOX21* and *SIX3*, accompanied by a specific decrease in H3K9me2, while global H3K9me2 at lamin-associated domains (LADs) remained unaffected (Fig. 2c-e, Ext. Data Fig. 2c)^21^. These results indicate that truncating ZNF462 mutations impair targeted recruitment of the G9A/GLP complex at ZNF462 targets, compromising localized heterochromatin formation and allowing premature activation of neuroectodermal gene programs.

**Fig. 2.**
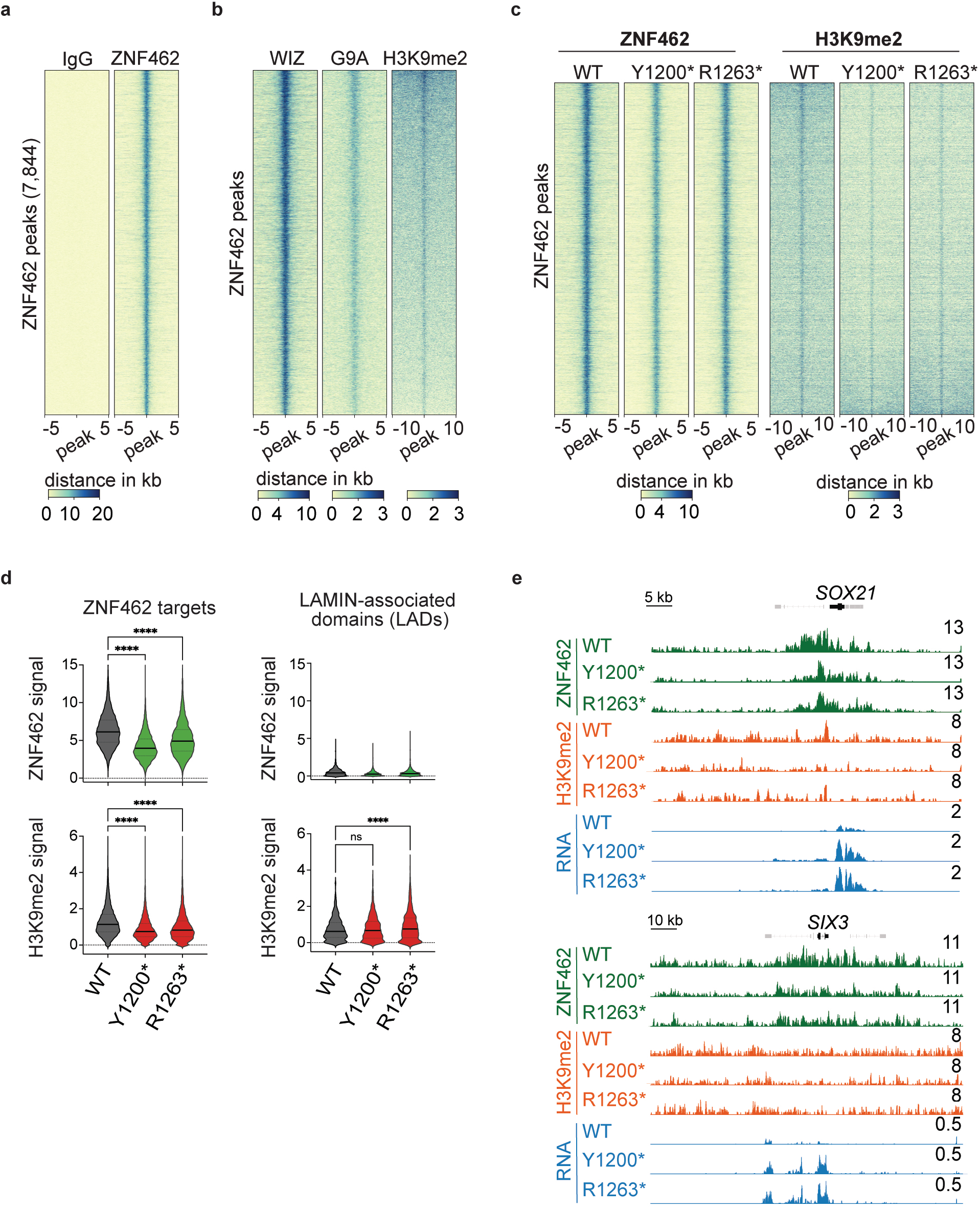
ZNF462 loss impairs targeting of G9A/GLP-dependent heterochromatin. **a)** Heatmaps of CUT&RUN signal of ZNF462 and matched IgG control at 7,844 high-confidence ZNF462-binding sites in WT hESCs. **b)** Heatmaps of WIZ, G9A, and H3K9me2 CUT&RUN signal at the same sites. **c)** Heatmaps showing ZNF462 and H3K9me2 CUT&RUN signal in WT, Y1200*, and R1263* hESCs at ZNF462 target sites. **d)** Violin plots showing ZNF462 and H3K9me2 signal at ZNF462 target sites and lamina-associated domains (LADs) in WT, Y1200*, and R1263* hESCs. Statistical significance was assessed using Friedman test followed by Dunn’s multiple comparisons test. ns, not significant; **** p < 0.0001. **e)** Genome browser tracks show ZNF462, H3K9me2 and mRNA levels at the *SOX21* and *SIX3* gene loci in WT, Y1200*, and R1263* hESCs.

### ZNF462 is enriched at primate-specific TE-derived enhancers

We previously showed that the ZNF462 mouse homolog, ZFP462, is enriched at evolutionarily young LTRs co-opted as functional enhancers in mESCs^18^. To determine whether TE targeting is conserved in human, we analyzed ZNF462 binding across repetitive DNA families in hESCs, revealing significant enrichment for LTR retrotransposons, specifically primate-specific ERV1 and ERVK classes including LTR7, LTR49, MER11C, and MER4B (Fig. 3a, b; Ext. Data Table 3). These evolutionarily young primate-specific LTRs are enriched for core pluripotency TF motifs (OCT4, SOX2, NANOG) and contribute to the hESC regulatory repertoire as active gene enhancers^22–24^. Consistent with this, motif enrichment analysis demonstrated strong enrichment for the composite OCT4-SOX2-TCF-NANOG recognition sequence (Fig. 3c, Ext. Data Table 4). Beyond pluripotency motifs, we identified significant enrichment for neurogenic TF motifs of the POU (BRN1, OCT6), SOX (SOX2, SOX3, SOX21), and ZIC (ZIC2, ZIC3) families. The unvalidated ZNF462 binding motif (JASPAR UN0616.1) as well as an uncharacterized motif (Unknown-ESC-element) containing the predicted ZNF462 core sequence (CAGCAGG) were also markedly enriched at ZNF462-bound regions. Notably, the CAGCAGG core is also contained within the ZIC family recognition sequences, suggesting that ZNF462 and ZIC factors potentially target a common DNA-binding motif (Fig. 3d). Integration of public ChIP-seq datasets confirmed that a subset of ZNF462 targets in hESCs is co-occupied by OCT4, SOX2, and ZIC2 (Fig. 3e, Ext. Data Fig. 3a), suggesting that ZNF462 antagonizes TF binding. Intersecting ZNF462 binding sites with SCREEN cCRE annotations revealed enrichment at distal enhancer-like elements, with ZNF462 targets displaying higher H3K4me1, H3K27ac, and ATAC-seq accessibility compared to matching unbound regulatory sequences (Fig. 3f, g)^25^. Thus, despite targeting species-specific LTRs, the fundamental mechanism of regulating young TE-derived enhancers is evolutionarily conserved.

**Fig. 3.**
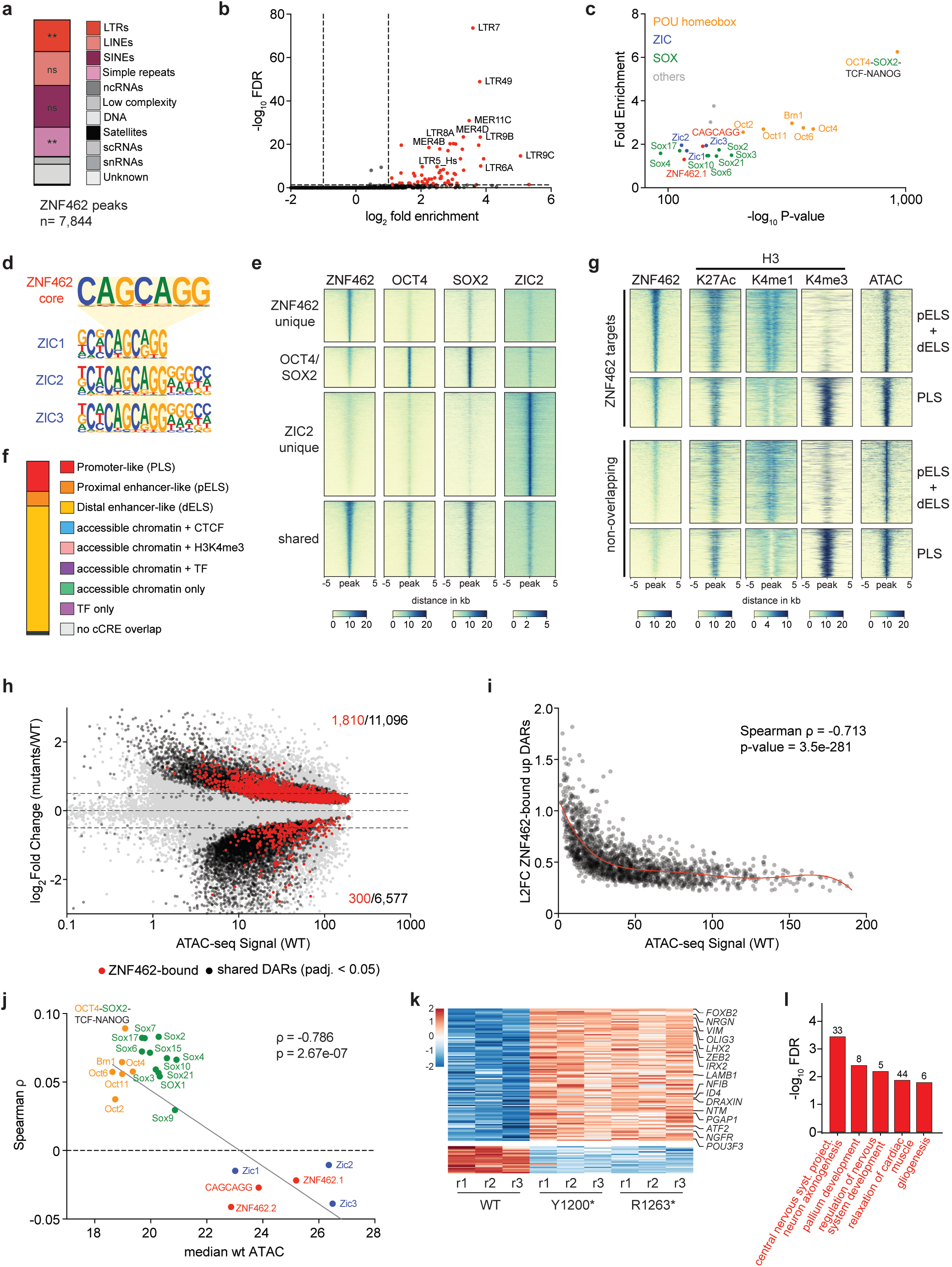
ZNF462 represses TE-enhancers enriched in POU, SOX and ZIC motifs in hESCs. **a)** Distribution of repeat-element classes overlapping ZNF462 target sites in hESCs. Statistical significance was assessed by one-sided Fisher exact test. ns, not significant; ** p < 0.01. **b)** Volcano plot showing fold enrichment and -log_10_ FDR of repeat-element subfamilies overlapping ZNF462 peaks. **c)** Scatter plot showing HOMER known-motif enrichment at ZNF462 target sites, plotted as fold enrichment versus -log_10_ P value. Transcription factor motif families are color-coded (POU, orange; ZIC, blue; SOX, green; other, grey); ZNF462-associated motifs are shown in red. **d)** Predicted ZNF462 and ZIC-family DNA-binding motifs showing similarity in their core sequence. **e)** Heatmaps of ZNF462, OCT4, SOX2 and ZIC2 binding signals at unique and overlapping targets. Shared indicates overlapping high-confidence binding sites between ZNF462, OCT4, SOX2 and ZIC2. **f)** Genomic annotation of ZNF462-bound regions based on SCREEN cCRE classifications. **g)** Heatmaps of ZNF462, histone modifications, and ATAC-seq signal at ZNF462-bound enhancer-like (pELS/dELS) and promoter-like (PLS) cCREs and matched cCREs lacking ZNF462 binding. **h)** Scatter plot showing ATAC-seq signal in WT hESCs vs. mean log_2_ fold change of ATAC-seq signal. Black dots indicate shared differentially accessible regions (DARs) in Y1200* and R1263* hESCs (FDR < 0.05). Red dots indicate shared ZNF462-bound DARs. **i)** Scatter plot of wild-type ATAC-seq signal vs mean log_2_ fold change of ZNF462-bound Shared-Up DARs (L2FC, mutants vs WT, y-axis). Red line shows median log_2_ fold change within each of 12 equal-width bins spanning the WT signal range; the shaded orange band indicates the interquartile range (25th–75th percentile) per bin. Spearman’s rank correlation is annotated. **j)** Scatter plot of median wild-type ATAC-seq signal at ZNF462-bound shared upregulated DARs overlapping with enriched TF motifs vs Spearman rank correlation (ρ) between motif presence and average log_2_ fold change (L2FC, mutant vs WT), quantifying the association between motif presence and accessibility gain. Dots are color-coded by TF family: POU (orange), SOX (green), ZIC (blue), and ZNF462-associated (red). The grey line shows a linear fit across all motifs; the motif-level Spearman correlation between baseline accessibility and ρ is annotated. **k)** Heatmap of RNA expression signal of DEGs linked to ZNF462-bound DARs (FDR < 0.05, |log_2_ fold change| ≥ 0.5). **l)** Gene Ontology biological process enrichment analysis of genes linked to ZNF462-bound upregulated DARs. Numbers above bars indicate fold enrichment. Significance is shown as -log_10_(FDR).

### ZNF462 represses DNA accessibility and TF binding at enhancers in hESCs

The enrichment of ZNF462 at active enhancers raised questions about its function as a repressor. To directly link ZNF462-dependent chromatin dynamics to transcriptional regulation, we performed ATAC-seq comparing wild-type and ZNF462 mutant hESCs, identifying 18,707 and 13,237 differentially accessible regions (DARs; FDR < 0.05) in the Y1200* and R1263* lines, respectively (Ext. Data Fig. 3b, c, Ext. Data Table 5). Chromatin accessibility changes from both mutant lines were highly correlated and overlapping. Intersecting DARs with ZNF462 occupancy confirmed its primary role as a repressor. The vast majority of ZNF462-bound DARs showed increased accessibility, with the most pronounced gains at ZNF462 targets with low baseline DNA accessibility in wild-type hESCs (Fig. 3h, i, Ext. Data Table 5).

Motif-accessibility correlation analysis revealed that POU and SOX family motifs were associated with low baseline DNA accessibility and strong positive correlations with accessibility gains upon ZNF462 loss (Fig. 3j, Ext. Data Fig. 3d). In contrast, ZIC- and the inferred ZNF462 (CAGCAGG)-containing sites exhibited high baseline DNA accessibility and no significant correlation with accessibility changes. To investigate these divergent chromatin responses, we controlled for baseline ATAC-seq signal to account for accessibility-driven motif enrichment across ZNF462 targets. Accessibility normalization (ATAC+GC content) exposed a substantial drop in ZIC (ZIC1-3) and ZNF462 (CAGCAGG) motif enrichment (Ext. Data Table 4), indicating that in hESCs ZNF462 occupancy at CAGCAGG/ZIC-motif containing enhancers is opportunistic and largely driven by pre-existing DNA accessibility, which aligns with the observation that ZNF462 loss elicits no local accessibility gains at these sites (Fig. 3j, Ext. Data Fig. 3d, e). In contrast, POU and SOX motifs remained highly enriched following ATAC-seq normalization, arguing that ZNF462 specifically targets and enforces G9A/GLP-mediated chromatin compaction at enhancers containing these motifs. Finally, linking ZNF462-bound enhancers to their nearest DEGs revealed that the vast majority were upregulated and enriched for nervous system development and neurogenesis (Fig. 3k, l). Together, these results demonstrate that ZNF462 haploinsufficiency increases DNA accessibility at early neuroectodermal enhancers containing POU and SOX motifs, driving aberrant gene activation and spontaneous neural differentiation via loss of G9A/GLP heterochromatin compaction.

### ZNF462 redirects to LTR5_Hs elements and lineage-specific enhancers

To investigate ZNF462 regulation during neuroectoderm development, we differentiated wild-type hESCs into NCCs and NPCs (Ext. Data Fig. 4a, b), modeling the developmental progenitors of tissues impacted in WKS. Flow cytometry analysis of cell-type specific markers CD271 (aka p75NTR, NGFR) and PAX6 confirmed generation of homogeneous NCC and NPC populations, respectively. Genome-wide CUT&RUN profiling revealed 6,822 ZNF462 peaks in NCCs and 4,411 in NPCs, with mostly distinct targets across NCCs, NPCs, and hESCs (Fig. 4a, Ext. Data Fig. 4c, Ext. Data Table 2). This cell-type-specific distribution was reflected in WIZ and H3K9me2 binding patterns, suggesting ZNF462 directs G9A/GLP to lineage-specific loci (Fig. 4b, c).

**Fig. 4.**
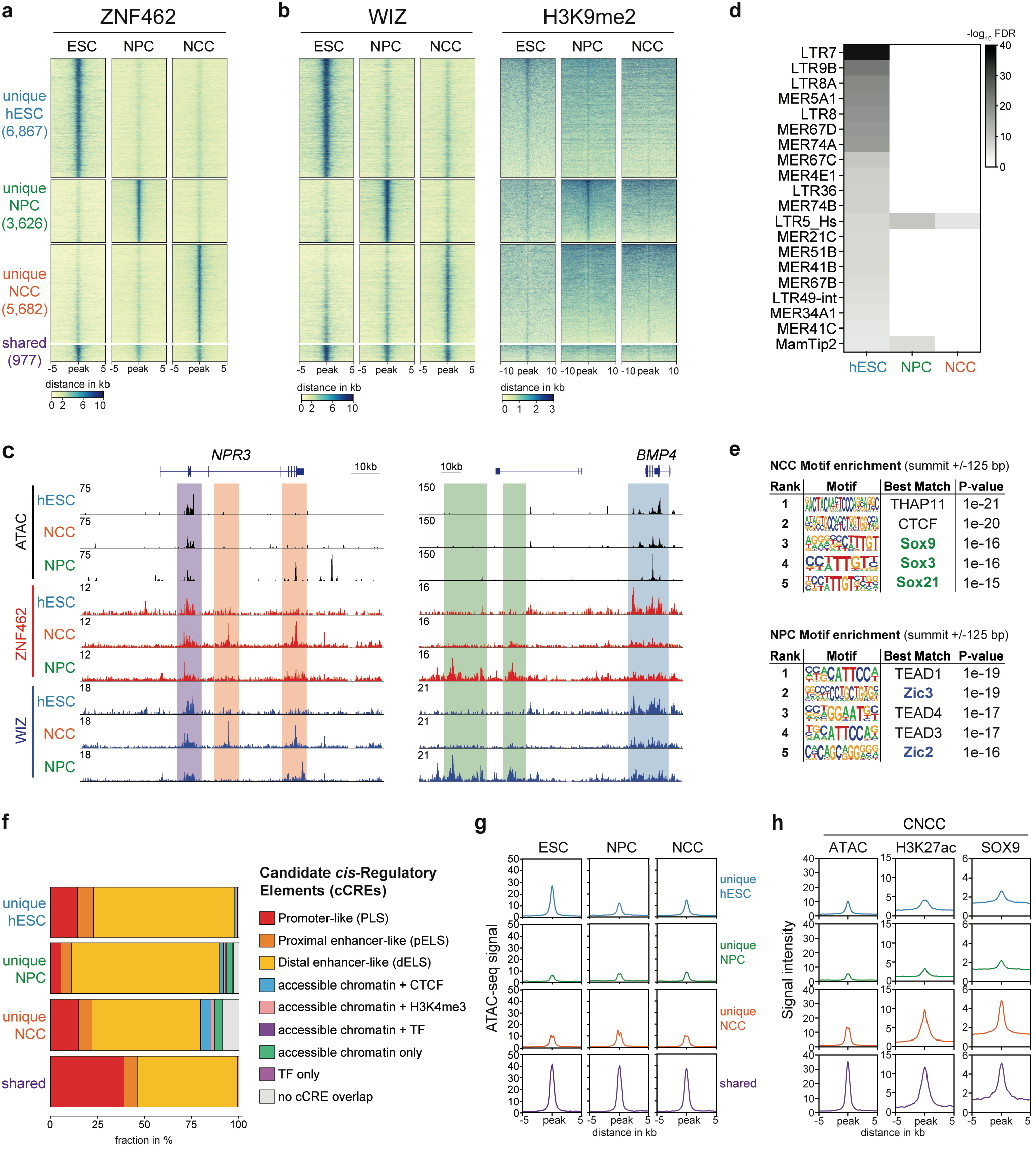
ZNF462 targets G9A/GLP to cell-type specific TE-enhancers. **a)** Heatmaps of ZNF462 CUT&RUN profiling reveal that high-confidence binding sites in hESCs, NPCs and NCCs are largely distinct. **b)** Corresponding heatmaps of WIZ and H3K9me2 enrichment at cell-type specific and shared clusters in hESCs, NPCs and NCCs. **c)** Genome browser tracks of ZNF462, WIZ and ATAC-seq signals at two representative gene loci, *NPR3* and *BMP4,* in hESC, NPCs and NCCs. Shaded regions highlight hESC-unique (blue), NPC-unique (green), NCC-unique (red) and shared (purple) ZNF462 targets. **d)** Heatmap showing significance of enrichment of repeat-element subfamilies at all hESC, NPC, and NCC ZNF462 targets. Significance was determined by False Discovery Rate (FDR). **e)** Top five known HOMER motifs at unique NCC and unique NPC ZNF462 targets using 250 bp summit-centered windows and a GC-matched genomic background. Motifs are ranked by HOMER hypergeometric P value. **f)** Distribution of SCREEN global GRCh38 cCRE classifications among unique hESC, unique NPC, unique NCC, and shared ZNF462 target sites. **g)** Metaplots of ATAC-seq signal at cell-type specific and shared ZNF462 target clusters in hESC, NPCs and NCCs. **h)** Metaplots of ATAC-seq, H3K27ac and SOX9 signals at cell-type specific and shared ZNF462 target clusters in CNCCs^29^.

Analyzing the TE distribution, we observed reduced ZNF462 binding in both NCCs and NPCs across most primate-specific LTR subfamilies (Fig. 4d, Ext. Data Fig. 4d, Ext. Data Table 3). However, ZNF462 peaks in both lineages remained enriched at LTR5_Hs elements of the HERV-K family (Fig. 4d, Ext. Data Fig. 4e). LTR5_Hs, the youngest hominoid-specific LTR subfamily, harbors both pluripotency motifs and binding sites for critical cranial NCC TFs, including TWIST1, SOX9/10, and TFAP2A, and contributes as functional enhancers in human craniofacial development^26–28^. Motif enrichment analysis of NCC-specific ZNF462 peaks revealed strong enrichment for core NCC regulators, including the SOX family members SOX9, SOX3 and SOX21 near the summit (+/− 125 bp), and SOX10, SOX2, SOX4 and SOX6 along with TFAP2A, TFAP2C and THAP11 (Ronin) within +/− 250 bp (Fig. 4e, Ext. Data Fig. 4f, Ext. Data Table 4). The predicted ZNF462 motif was not enriched at NCC-specific sites, suggesting that ZNF462 recruitment to NCC enhancers occurs via alternative sequence motifs or co-factor interactions. Conversely, NPC-specific peaks exhibited enrichment of both ZNF462 and ZIC2/3 binding sites, yet without SOX family motifs among top ranks (Fig. 4e, Ext. Data Fig. 4f). In addition, NPC peaks showed selective enrichment for neurogenic TFs including TEAD family (TEAD1, TEAD2, TEAD3, TEAD4) and Otx2.

Controlling for local chromatin accessibility using ATAC-seq-matched backgrounds confirmed this inverse pattern of NPC peaks being significantly enriched for CAGCAGG and ZIC factor motifs, whereas NCC peaks displayed CAGCAGG depletion while retaining selective SOX family enrichment (Ext. Data Fig. 4g, Ext. Data Table 4). Together, these findings demonstrate that ZNF462 is dynamically redirected from pluripotent LTRs to hominoid-specific LTR5_Hs elements and lineage-specific TF binding motifs, establishing cell-type-specific regulatory frameworks of repression at ZIC-driven enhancers in NPCs versus SOX-driven enhancers in NCCs.

### ZNF462-bound lineage-specific enhancers are inaccessible

To characterize lineage-specific binding sites, we intersected NCC and NPC ZNF462 targets with SCREEN cCRE annotations, confirming enrichment at distal enhancer-like signatures (Fig. 4f). ATAC-seq in NPCs and integrated NCC datasets revealed that, unlike hESC targets with high baseline DNA accessibility, lineage-specific targets in NCCs and NPCs exhibited remarkably low DNA accessibility in progenitor states (Fig. 4g, Ext. Data Fig. 3b, 4h). We hypothesized that ZNF462 actively represses these enhancers to prevent untimely activation. Analyzing published datasets, we found that NCC ZNF462 targets displayed marked gains in DNA accessibility, H3K27ac deposition, and SOX9 recruitment in committed cranial neural crest cells (CNCCs)^29^, as well as SOX10 binding in transformed melanocytes^30^ (Fig. 4h, Ext. Data Fig. 4i, j). This suggests that ZNF462-bound progenitor targets mature into functional, active enhancers upon lineage progression. These findings argue that ZNF462 acts as an epigenetic gatekeeper that represses DNA accessibility at lineage-specific enhancers in progenitor cells to ensure proper developmental timing.

### ZNF462 loss aberrantly accelerates cranial neural crest specification

To model the functional impact of ZNF462 loss on neural cell types affected in WKS, we differentiated wild-type and mutant hESCs into NCCs and NPCs. During NCC differentiation, flow cytometry for CD271 revealed a striking temporal shift. ZNF462 mutant cells exhibited significantly more rapid and uniform CD271 induction by day five, yielding over 70% CD271-positive cells compared to only 25% in wild-type (Fig. 5a, b). Wild-type cells required an additional three days to reach comparable levels, indicating that ZNF462 restricts the tempo of NCC specification (Ext. Data Fig. 4a). Transcriptome analysis at day five identified 1,600 and 1,921 DEGs (FDR < 0.05, |L2FC| ≥ 1) in Y1200* and R1263* mutants, respectively (Ext. Data Fig. 5a, b, Ext. Data Table 1). R1263* exhibited more than twice as many unique downregulated DEGs as Y1200* (506 vs. 204), reflecting more pronounced disruption (Ext. Data Fig. 5c). Both lines showed significant downregulation of early neural plate border specifiers (*MSX1, ZIC2, ZIC3, ZIC4, ZIC5*) and upstream signaling ligands (*WNT1, WNT3, WNT5B, WNT9A, WNT9B, FGF11*), alongside marked upregulation of definitive NCC specifiers and EMT drivers (*SOX10, SNAI2, TFAP2A, TFAP2B, SOX8, LMO4*) (Fig. 5C, D). R1263* mutants uniquely downregulated additional border regulators (*ZIC1, MSX2, WNT3A, RSPO3, BMP4, GDF6*) while exclusively upregulating ectomesenchymal drivers (*ETS1, PDGFRA*), suggesting the transition toward peripheral glial identities (*SOX8/10* high, *SOX9* low) (Fig. 5d).

**Fig. 5.**
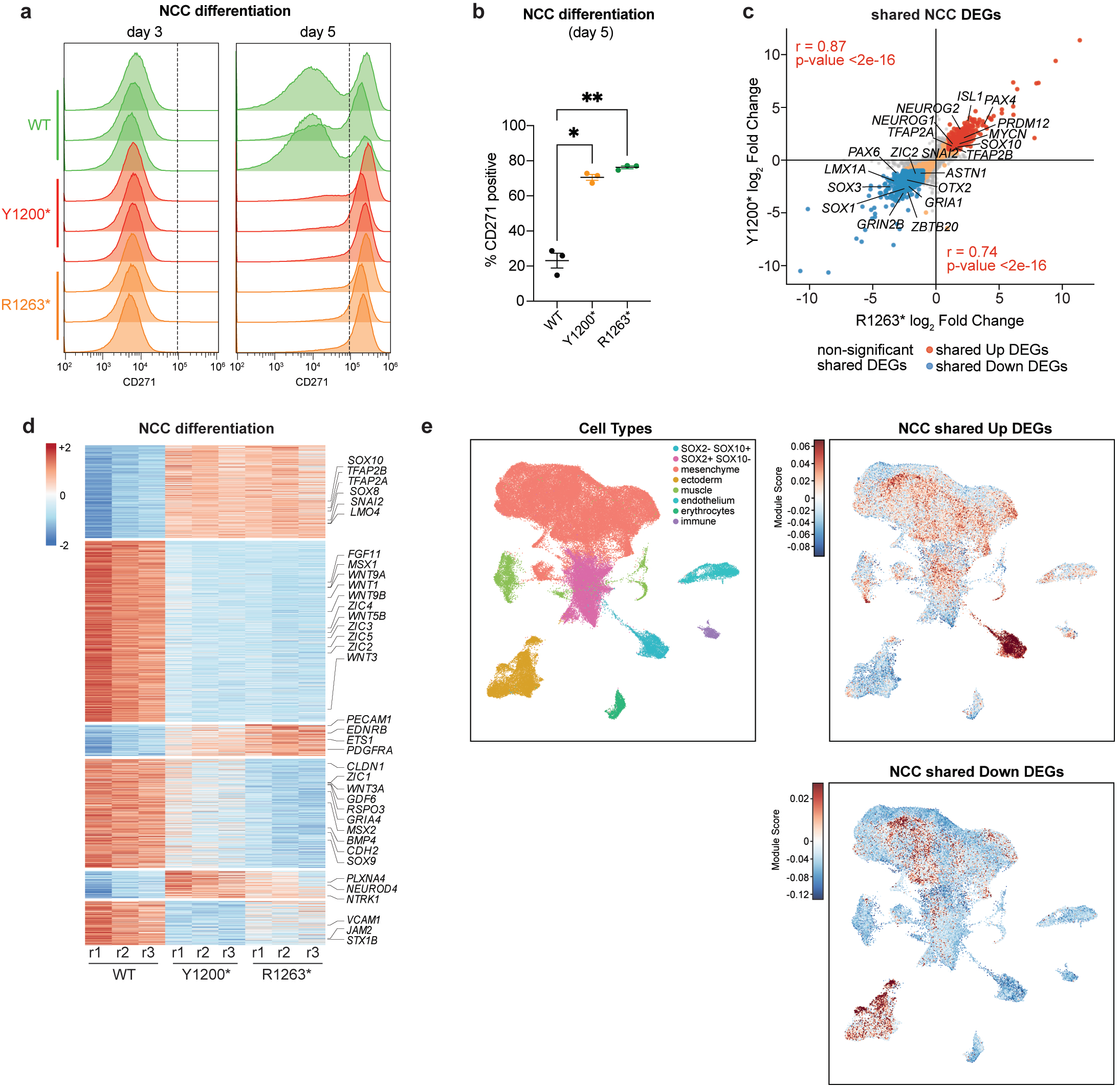
ZNF462 restricts timing of neural crest cell lineage specification. **a)** Representative flow cytometry histograms of CD271 expression during *in vitro* NCC differentiation of WT, Y1200*, and R1263* hESCs at days 3 and 5. **b)** Quantification of the fraction of CD271-positive cells at day 5 of NCC differentiation. Bars represent mean ± SEM from n = 3 independent matched differentiation experiments. Statistical significance was assessed by one-way repeated-measures ANOVA with Geisser–Greenhouse correction followed by Dunnett’s multiple comparisons test using WT as the reference group. *p < 0.05; **p < 0.01. **c)** Pairwise DEG (FDR < 0.05 and |log_2_ fold change| ≥ 1) comparison of R1263* and Y1200* mutant cells relative to WT at 5 days of NCC differentiation. Shared upregulated DEGs highlighted in red, and shared downregulated DEGs highlighted in blue. Pearson correlation coefficients and corresponding p-values are shown. **d)** Heatmap of shared mutant DEGs showing RNA expression in WT, Y1200*, and R1263* cells at 5 days of NCC differentiation. Highlighted are critical regulators of neural plate border and NCC specification. **E)** UMAP projection of 95,892 cells from a human craniofacial single-cell atlas^31^, colored by cell type (left) or by module score for the NCC shared-up (top right) and NCC shared-down (bottom right) differentiation signatures derived from ZNF462 mutant versus wild-type NCC bulk RNA-seq. Cell types: cranial neural crest cells (CNCC, SOX2⁻/SOX10⁺, n = 3,294), neural progenitors (SOX2⁺/SOX10⁻, n = 14,412), mesenchyme (n = 58,417), ectoderm (n = 9,652), muscle (n = 4,212), endothelium (n = 3,424), erythrocytes (n = 1,609), and immune (n = 872). **f)** Gene Ontology (GO) analysis of shared NCC DEGs. Red bars indicate GO-terms of shared upregulated DEGs, blue bars indicate GO-terms of shared downregulated DEGs.

To confirm that these changes reflect a coherent acceleration along physiological trajectories, we mapped DEG signatures onto single-cell RNA-seq (scRNA-seq) atlases of human embryonic/gastruloids and cranial NCC subtypes^31^. In the embryonic/gastruloid atlas (CS12-CS20), the NCC-up DEG signature was specifically enriched in *SOX2⁻/SOX10⁺* CNCCs and depleted in neural progenitors (*SOX2⁺/SOX10⁻*) and ectoderm (Fig. 5e; Ext. Data Fig. 5d). Stage-stratified analysis confirmed positive scores in CNCCs across all Carnegie stages versus negative scores in progenitors (Ext. Data Fig. 5e, f). Trajectory analysis across 11 CNCC subtypes revealed a significant gradient, with highest NCC-up DEG scores in early committed CNCCs and lower scores in mature derivatives. The NCC-down DEG signature showed the inverse pattern. These results argue that ZNF462 mutant cells display accelerated differentiation toward early CNCC identity. By failing to repress enhancers enriched in SOX family and other core NCC motifs, ZNF462 haploinsufficiency drives an abnormally rapid transition into the cranial neural crest program, establishing ZNF462 as an essential gatekeeper that restricts premature specification and dictates human developmental tempo.

### ZNF462 haploinsufficiency stalls cortical neurogenesis without impairing neurogenic potential

While ZNF462 loss accelerated NCC specification, its impact on NPC differentiation was morphologically subtle. PAX6 immunostaining at day 27 revealed homogeneous neural progenitor populations with no significant difference in PAX6 positivity between wild-type and mutant lines (Fig. 6a, Ext. Data Fig. 6a). However, transcriptome profiling identified 676 and 575 DEGs (FDR < 0.05, |L2FC| ≥ 0.5) in Y1200* and R1263* NPCs, respectively, with largely overlapping and highly correlated changes between mutant clones (Fig. 6b; Ext. Data Fig. 6a-c; Ext. Data Table 1). Shared DEGs revealed aberrant activation of early signaling modulators and multipotent regulators (Fig. 6c). Mutant NPCs upregulated WNT and BMP components (*SMAD9, TCF7, AXIN2, LEF1, GDF7, ID1, BMP4*), early regional markers (*FOXG1, NFIA*), and neural plate border specifiers (*ZIC2, ZIC5*). This was accompanied by downregulation of synaptic receptors (*GRIN2A, GABRB2, SYT2, SV2A*) and axon guidance drivers (*PLXNA2, DISC1, FLRT2, DLG2, BRINP1*), suggesting a developmental delay.

**Fig. 6.**
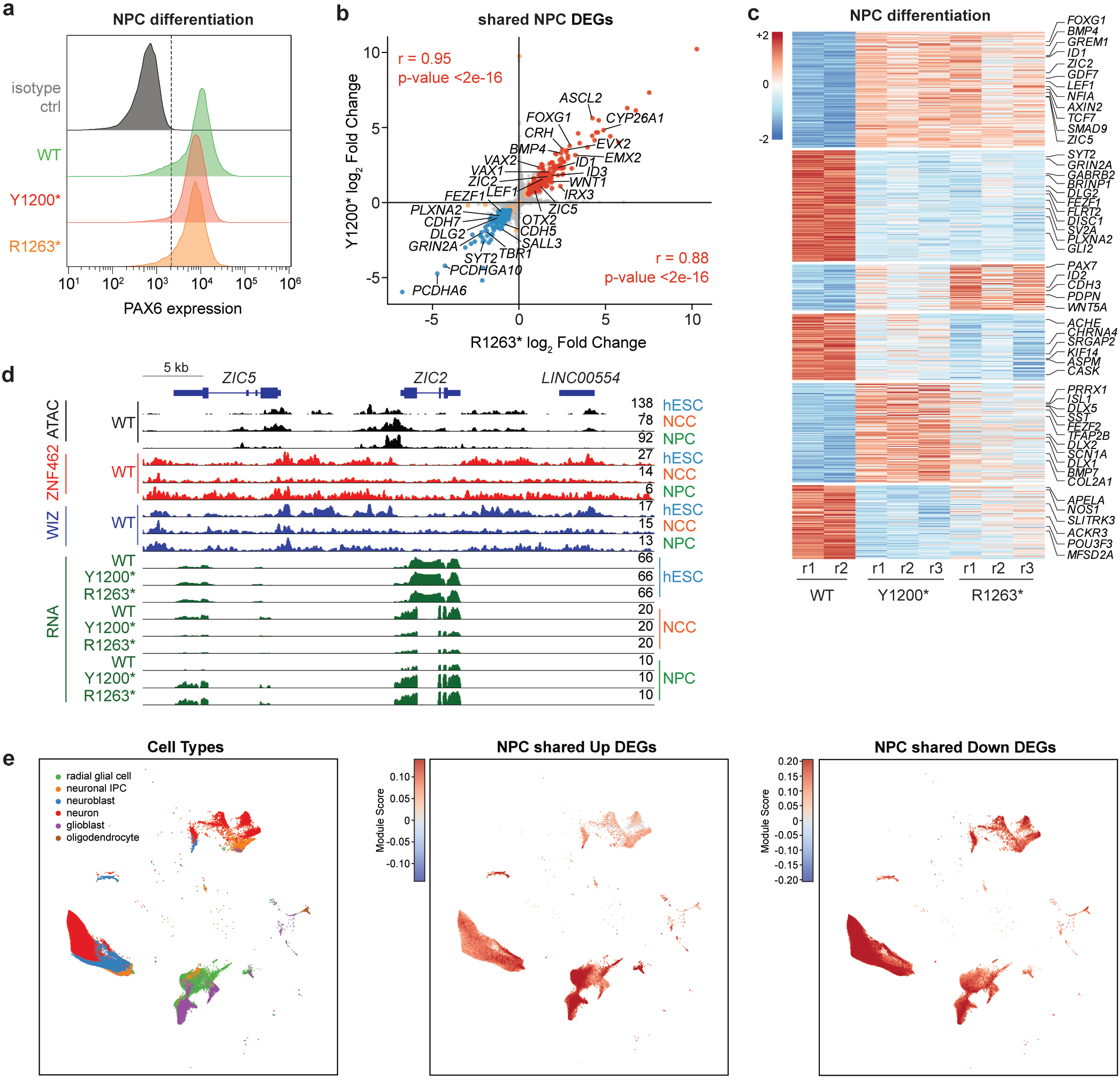
ZNF462 is required for proper timing of neuronal cell lineage specification. **a)** Representative flow cytometry histograms showing PAX6 expression after 27 days of NPC differentiation of WT, Y1200*, and R1263* hESC. **b)** Pairwise DEG (padj. < 0.05 and |log_2_ fold change| ≥ 0.5) comparison of R1263* and Y1200* mutant cells relative to WT at 27 days of NPC differentiation. Shared upregulated DEGs highlighted as red, and shared downregulated DEGs highlighted as blue. Pearson correlation coefficients and corresponding p-values are shown. **c)** Heatmap of shared mutant DEGs showing RNA expression in WT, Y1200*, and R1263* cells at 27 days of NPC differentiation. Highlighted are critical regulators of neural plate border and neuronal specification. **D)** UMAP projection of 49,997 cells from first-trimester human telencephalon single-cell atlas^33^, colored by cell type (left) or by module score for the NPC shared-up (top right) and NPC shared-down (bottom left) differentiation signatures derived from ZNF462 mutant versus wild-type NPC bulk RNA-seq. Cell type reference: radial glia (n = 16,427), neuron (n = 16,804), neuroblast (sensu Vertebrata) (n = 8,387), neuroblast (sensu Nematoda and Protostomia) (n = 4,301), glioblast (n = 3,950), and oligodendrocyte (n = 128). **E)** Genome browser tracks showing ATAC-seq, ZNF462 and WIZ CUT&RUN, and RNA-seq signal at the *ZIC2* gene locus in hESCs, NCCs, and NPCs. RNA-seq tracks show WT, Y1200*, and R1263* cells for each lineage.

Mechanistically, we found that ZNF462 functions as a critical repressor of the neural plate border program not only by restricting ZIC factor activity at downstream border enhancers but also by directly binding and repressing the *ZIC2* gene locus itself, which is markedly upregulated in *ZNF462* mutant NPCs (Fig. 6d). Because ZIC2 governs early neural border patterning and sustains signaling cascades that prevent neurogenesis, and ZIC factor overexpression is known to prevent neuronal differentiation *in vivo*^14,32^, loss of ZNF462 repression maintains expression and activity of border specifiers.

To confirm this reflects coherent deceleration, we mapped NPC DEG signatures onto a 49,997-cell scRNA-seq atlas of the developing human telencephalon (weeks 9–15)^33^. The NPC-up DEG signature exhibited a significant downward gradient along the cortical differentiation trajectory, peaking in radial glial cells and progressively decreasing through neuroblasts to mature neurons (Fig. 6e; Ext. Data Fig. 6e). Conversely, the NPC-down DEG signature displayed the inverse pattern, with the lowest scores in radial glial cells and highest in neuroblasts and neurons. These results suggest that ZNF462 loss traps NPCs in an early radial glial progenitor state by failing to silence *ZIC2* and the neural plate border regulatory networks that normally inhibit neurogenesis^14,32^.

To determine whether this radial glial retention reflects a permanent lineage block or an altered developmental rate, we subjected wild-type and *ZNF462* mutant NPCs to extended cortical neuron differentiation. Following prolonged culture, *ZNF462* mutant NPCs successfully generated MAP2-positive cortical neurons, demonstrating that their neurogenic capacity remains fully intact (Ext. Data Fig. 6f). At this same time point, Fluo-4 calcium imaging detected spontaneous calcium transients in both mutant and wild-type control neurons, confirming that mutant cells acquire functional activity without an overt defect in electrophysiological maturation (Ext. Data Fig. 6g). This preservation of terminal neuronal differentiation and spontaneous calcium activity demonstrates that ZNF462 haploinsufficiency does not abolish intrinsic neurogenic potential. Instead, loss of ZNF462 establishes a transient developmental barrier that decelerates progenitor exit and cortical maturation. Together, these functional data provide compelling evidence that ZNF462 operates as a lineage-specific epigenetic gatekeeper of human developmental tempo rather than a determinant of lineage competence.

## Discussion

The neural plate border represents a pivotal developmental junction where central nervous system precursors and neural crest lineages diverge from a shared ectodermal progenitor pool. Precise temporal control of lineage exit at this border is essential to synchronize the co-development of the skull and brain^1,7,10,13,16^. Our findings indicate that ZNF462 functions as an essential epigenetic gatekeeper coordinating the timing of this bifurcation. Rather than enforcing a static transcriptional state, ZNF462 exerts stage-specific, lineage-dependent repression across both lineages, ensuring that cortical neuronal progenitors proceed toward differentiation on schedule while restraining neural crest specification until the appropriate developmental window. We speculate that uncoupling this synchronized control by accelerating neural crest specification while decelerating neural progenitor exit disrupts the coordinated assembly of the head. Because mutant progenitors retain full intrinsic potential to form functional neurons and neural crest derivatives, WKS pathogenesis represents a primary defect in developmental tempo rather than lineage specification, providing a novel conceptual framework for the co-occurring neurodevelopmental delays and craniofacial dysmorphisms in WKS.

A central question is how ZNF462 selects its genomic targets across distinct developmental states. In hESCs, although the inferred CAGCAGG consensus is enriched, top-ranked motifs correspond to POU and SOX family TF recognition sequences. In contrast, the enrichment of the shared ZNF462/ZIC motif is lost at ZNF462 peaks after ATAC-seq normalization, suggesting that targeting in pluripotency is selective for LTRs with POU and SOX family motifs, and that CAGCAGG motif binding is largely opportunistic, driven by the high baseline DNA accessibility. Upon neural lineage commitment, ZNF462 undergoes a dramatic targeting shift to hominoid-specific LTR5_Hs elements and lineage-specific enhancers that are relatively inaccessible in precursor states. Because these targets lack open chromatin, recruitment cannot be explained by passive accessibility alone. The lineage-specific motif enrichment of ZNF462-bound enhancers suggests lineage-divergent targeting principles. In NCCs, SOX family motifs are the strongest accessibility-independent features of ZNF462 binding, suggesting that SOX-driven enhancers serve as primary targets of ZNF462 recruitment. In NPCs, ZIC motif enrichment survives accessibility normalization while SOX family motifs are absent, indicating that ZNF462 may exhibit context-dependent selectivity for ZIC-containing regulatory elements. While context-dependent deployment of C_2_H_2_ arrays is documented in other factors (such as ZFP335, CTCF, and Kaiso)^34–36^, future structural and biochemical studies will be needed to determine whether ZNF462 utilizes distinct zinc finger subsets or co-factor interactions across these cell types.

Crucially, the specific target enhancer program correlates with the direction of the temporal shift upon ZNF462 loss. In wild-type NPCs, ZNF462 enforces repression of the *ZIC2* gene locus and ZIC-associated border enhancers, silencing early border specifiers (*ZIC, WNT, BMP*) to allow progenitors to commit to cortical neurogenesis on schedule. Upon ZNF462 loss, persistent expression of *ZIC2* and border specifiers creates a developmental barrier that traps NPCs in a radial glial state and decelerates neurogenesis^14,32^. Importantly, our extended differentiation experiments demonstrate that this progenitor stall does not prevent eventual neuronal maturation or functional network assembly, confirming that ZNF462 controls developmental kinetics rather than lineage potential. Conversely, in NCCs, ZNF462 represses SOX-driven enhancers. Loss of ZNF462 leads to premature activation of definitive crest specifiers (*SOX10, SNAI2, TFAP2A/B*), accelerating EMT. Single-cell trajectory mapping confirms that both transcriptional shifts represent coherent movements along physiological developmental pathways, supporting the notion that haploinsufficiency of a single epigenetic regulator can produce opposing kinetic shifts in two lineages arising from the same progenitor territory.

Our analysis of clinical variants provides a molecular rationale for phenotypic variability in WKS. While both Y1200* and R1263* produce overlapping DEG profiles and neurogenic stalling in NPCs, they diverge significantly in NCCs, with R1263* inducing more severe transcriptional disruption, heightened NCC acceleration, and premature EMT, correlating with the more severe craniofacial dysmorphism observed in R1263* patients^3,4^.

The dynamic redirection of ZNF462 to LTR5_Hs elements in differentiated cells connects retrotransposon evolution to human developmental timing. As the youngest hominoid-specific LTR subfamily, LTR5_Hs represents a recently acquired enhancer platform for cranial NCC TFs^26–28^. Ultimately, our findings suggest that developmental timing disorders represent a distinct mechanistic category of human congenital malformations. WKS is not a defect of cell fate specification or differentiation competence, as mutant NPCs retain full capacity to generate functional cortical neurons upon extended culture, but rather a disorder of temporal coordination, where the pace of lineage commitment is asynchronously altered in opposite directions. This concept provides a generalizable framework for understanding other pleiotropic neurodevelopmental syndromes caused by haploinsufficiency of chromatin regulators controlling developmental timing.

## Supporting information

Ext. Data Table 1

Ext. Data Table 2

Ext. Data Table 3

Ext. Data Table 4

Ext. Data Table 5

Ext. Data Table 6

Ext. Data Table 7

Ext. Data Table 8

## Acknowledgements

We are grateful to all members of the Bell and Dou laboratories, Unmesh Jadhav and Matthew Deardorff for support, feedback and discussions. We thank Sayyada Shahid, Joanna Wysocka, Javeed Mohammed, and Alireza Lorzadeh for experimental advice and helpful discussions. We thank Gage Crump, Kritika Kasliwal and Matthias Stadtfeld for feedback on the manuscript. We apologize to colleagues whose work could not be cited due to space limitations. Funding: O. B. was supported by the National Institute of Mental Health (R01MH122565) and the Norris Comprehensive Cancer Center of USC and its NCI Award (P30CA014089). R. Y. was supported by EMBO Long-Term Fellowship (ALTF 256-2015). M. B. was supported by the National Institutes of Health (NIDDK K99DK143293). G. Q. was supported by the National Institute of Mental Health (R01MH13651 and R01MH138371). The funders had no role in study design, data collection and analysis, decision to publish or preparation of the manuscript.

## Author Contributions Statement

E. A. and O. B. initiated and designed the study. R. Y. generated the mutant hESC lines. E.A. performed the differentiation of hESCs, the genomic and transcriptomic analysis of hESC, NPCs and NCCs, including data analysis. S. M. contributed to the differentiation and genomic profiling of NCCs. C. S. and S. R. performed parts of the genomic data analysis and provided advice. E. A. and O. B. analyzed transcriptome and epigenome data. M. B. performed the cortical neuron differentiation and analysis. O. B. supervised all aspects of the project. The manuscript was prepared by E. A. and O. B.. G. Q., M. B. and S. M. edited the manuscript. All authors discussed results and commented on the manuscript.

---

**Ext. Data Fig 1.**
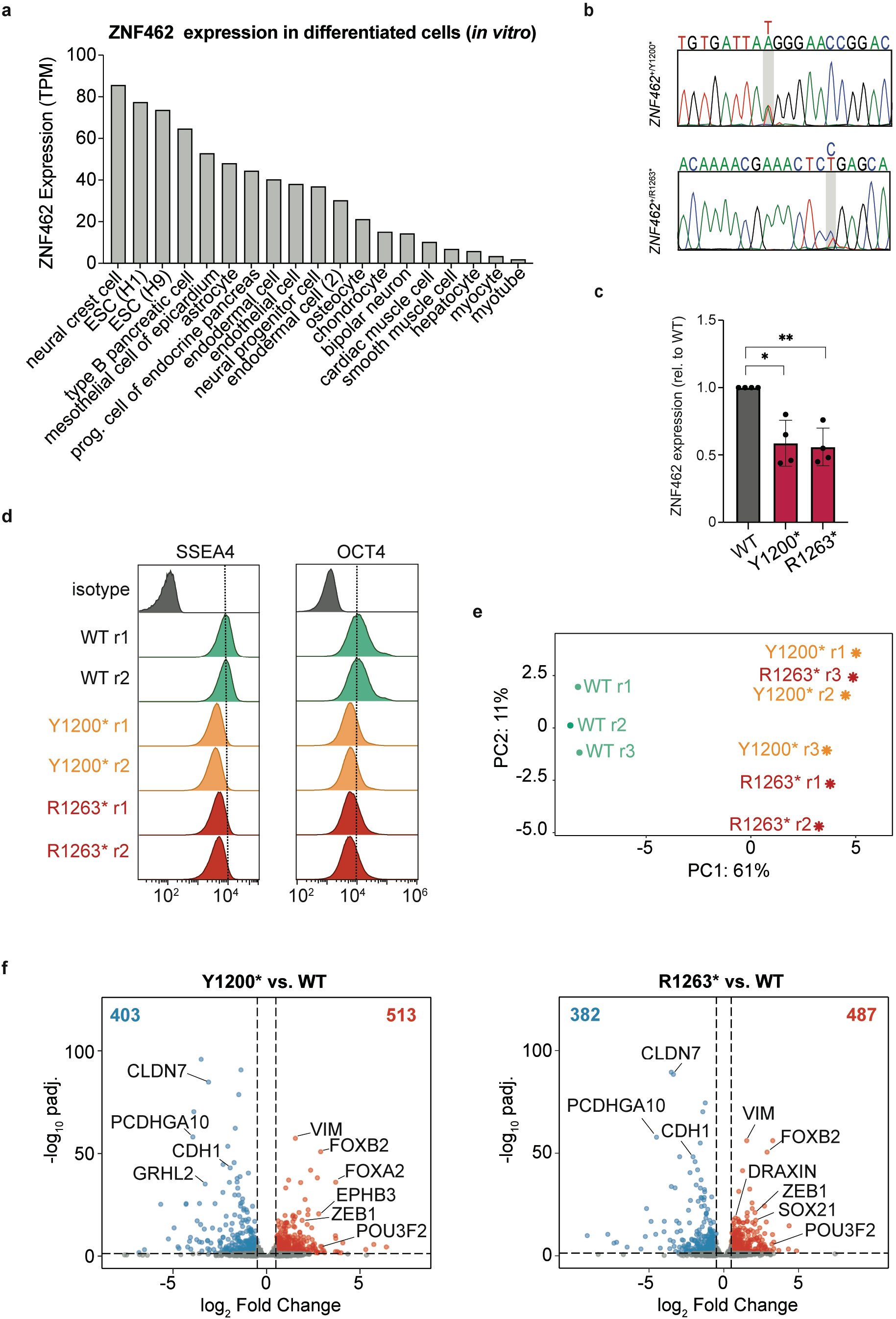
ZNF462 is required for silencing of neuroectodermal genes in hESC. **a)** Bar plot of ZNF462 RNA expression across *in vitro* differentiated human embryonic stem cell-derived cell types (ENCODE)^25^. **b)** Sanger sequencing chromatograms confirming heterozygous sequence variants in ZNF462 mutant hESC lines Y1200* and R1263*. **c)** Quantification of ZNF462 protein abundance in WT, Y1200*, and R1263* hESCs relative to WT. Bars represent mean ± SEM from n = 4 independent matched experiments. Statistical significance was assessed by one-way repeated-measures ANOVA with Greenhouse-Geisser correction followed by Dunnett’s multiple-comparisons test using WT as the reference group. * P < 0.05; ** P < 0.01. **d)** Representative flow cytometry histograms showing SSEA4 and OCT3/4 expression in WT, Y1200*, and R1263* hESCs. Isotype controls are shown in gray. **e)** Principal component analysis (PCA) of bulk RNA-seq data from WT, Y1200*, and R1263* hESCs. **f)** Volcano plots showing differential gene expression in Y1200* and R1263* hESCs relative to WT. Upregulated and downregulated DEGs are highlighted in red and blue, respectively. Dashed lines indicate the FDR and log2 fold-change thresholds used for differential expression analysis. Numbers indicate the total numbers of upregulated and downregulated DEGs (FDR < 0.05, |log2 fold change| ≥ 0.5).

**Ext. Data Fig. 2.**
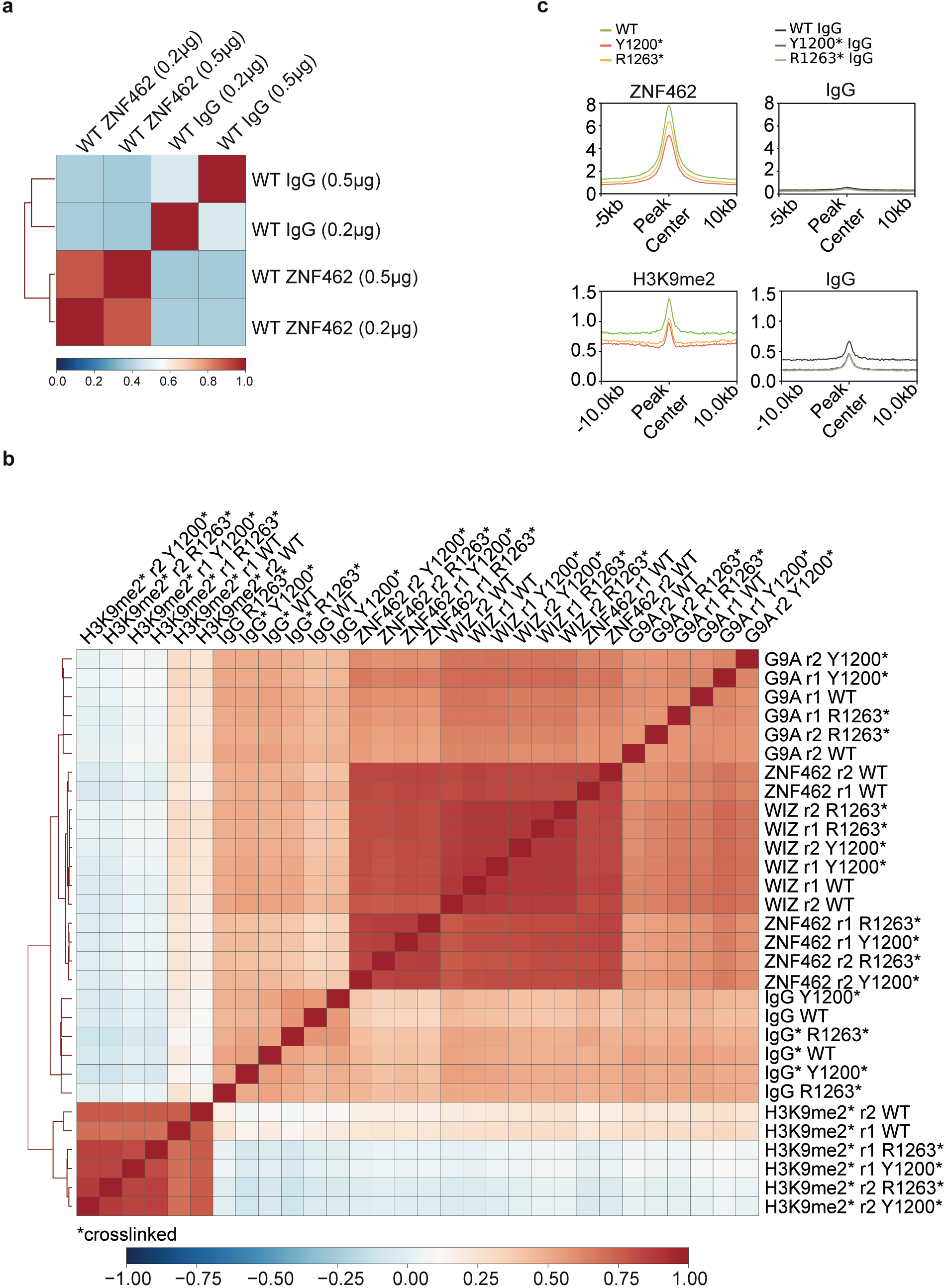
ZNF462 loss impairs targeting of G9A/GLP-dependent heterochromatin. **a)** Pearson correlation matrix comparing ZNF462 CUT&RUN and matched IgG signal across WT high-confidence ZNF462 peak regions using different amounts of ZNF462 antibody and corresponding IgG controls. **b)** Pearson correlation matrix comparing genome-wide CUT&RUN signal across ZNF462, WIZ, G9A, H3K9me2, and IgG samples in WT, Y1200*, and R1263* hESCs. Crosslinked samples are indicated by an asterisk (*). **c)** Metaplots showing ZNF462 and H3K9me2 CUT&RUN signal in WT, Y1200*, and R1263* hESCs centered on WT high-confidence ZNF462 peak regions. Corresponding IgG metaplots are shown as controls.

**Ext. Data Fig. 3.**
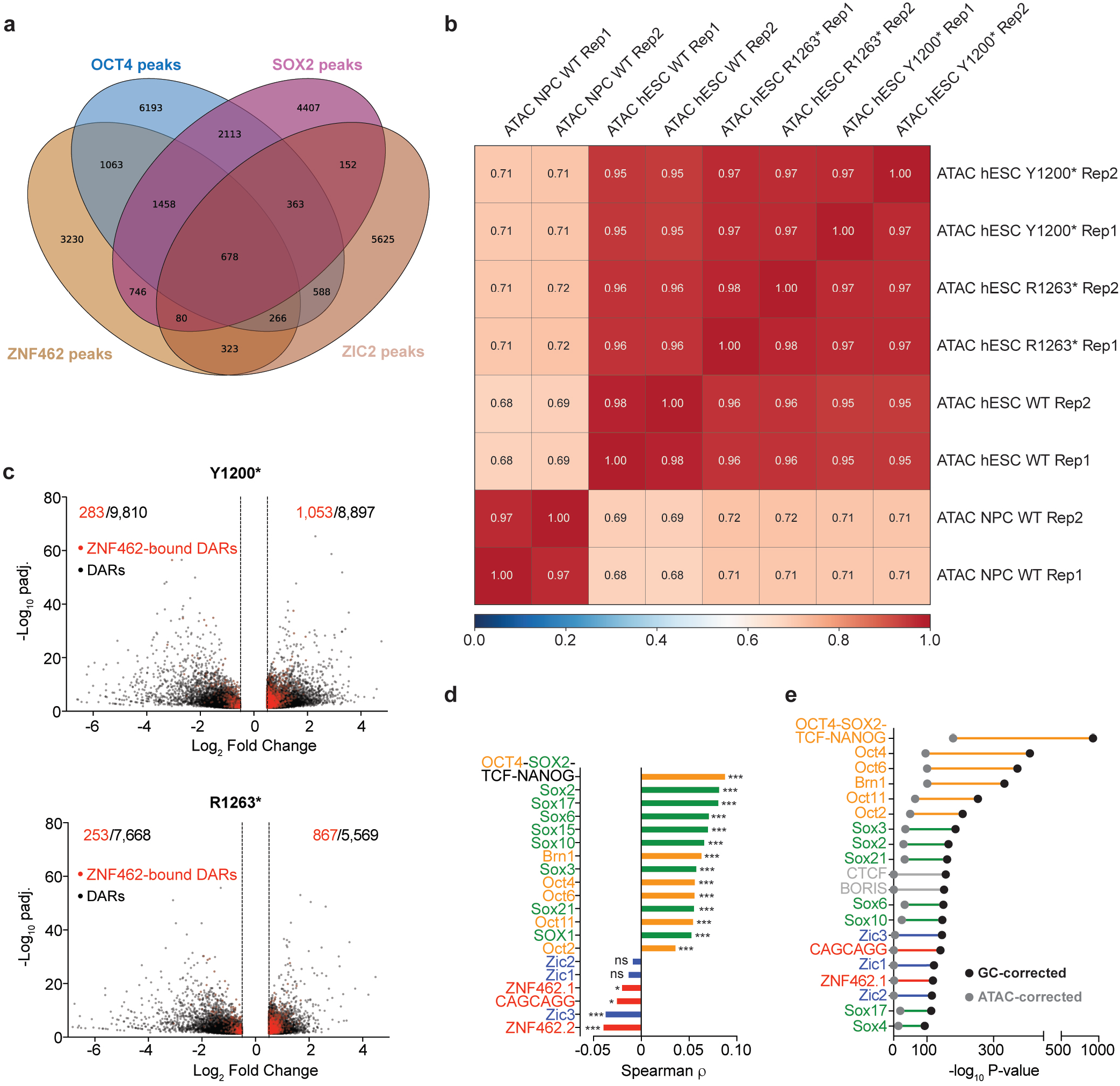
ZNF462 represses TE-enhancers enriched in POU and SOX motifs in hESCs. **a)** Venn diagram showing overlap among high-confidence ZNF462, OCT4, SOX2, and ZIC2 binding sites in hESCs. **b)** Pearson correlation matrix of ATAC-seq signal across WT, Y1200*, and R1263* hESCs and WT neural progenitor cells (NPCs). **c)** Volcano plots showing changes in chromatin accessibility in Y1200* and R1263* hESCs relative to WT. Black dots and numbers indicate differentially accessible regions (DARs; FDR < 0.05 and |log2 fold change| ≥ 0.5), and red dots and numbers indicate the subset of DARs overlapping ZNF462 binding sites. **d)** Bar plot showing Spearman rank correlation (ρ) between the binary presence of the selected TF family motifs and mean ATAC-seq log_2_ fold change (mutant vs WT) across 1,810 ZNF462-bound shared-Up DARs. Motifs are colored by family: POU (orange), SOX (green), ZIC (blue), and ZNF462 (red). Asterisks indicate significance after Benjamini–Hochberg correction, ns, not significant. * p < 0.05, *** p < 0.001. Positive Spearman ρ indicates that motif presence is associated with greater accessibility gain following ZNF462 loss. **e)** Comparison of HOMER motif enrichment at ZNF462 target sites using GC-matched genomic background regions (black) or regions jointly matched for GC content and WT ATAC-seq signal (gray). Motifs are colored by family as in **d**).

**Ext. Data Fig. 4.**
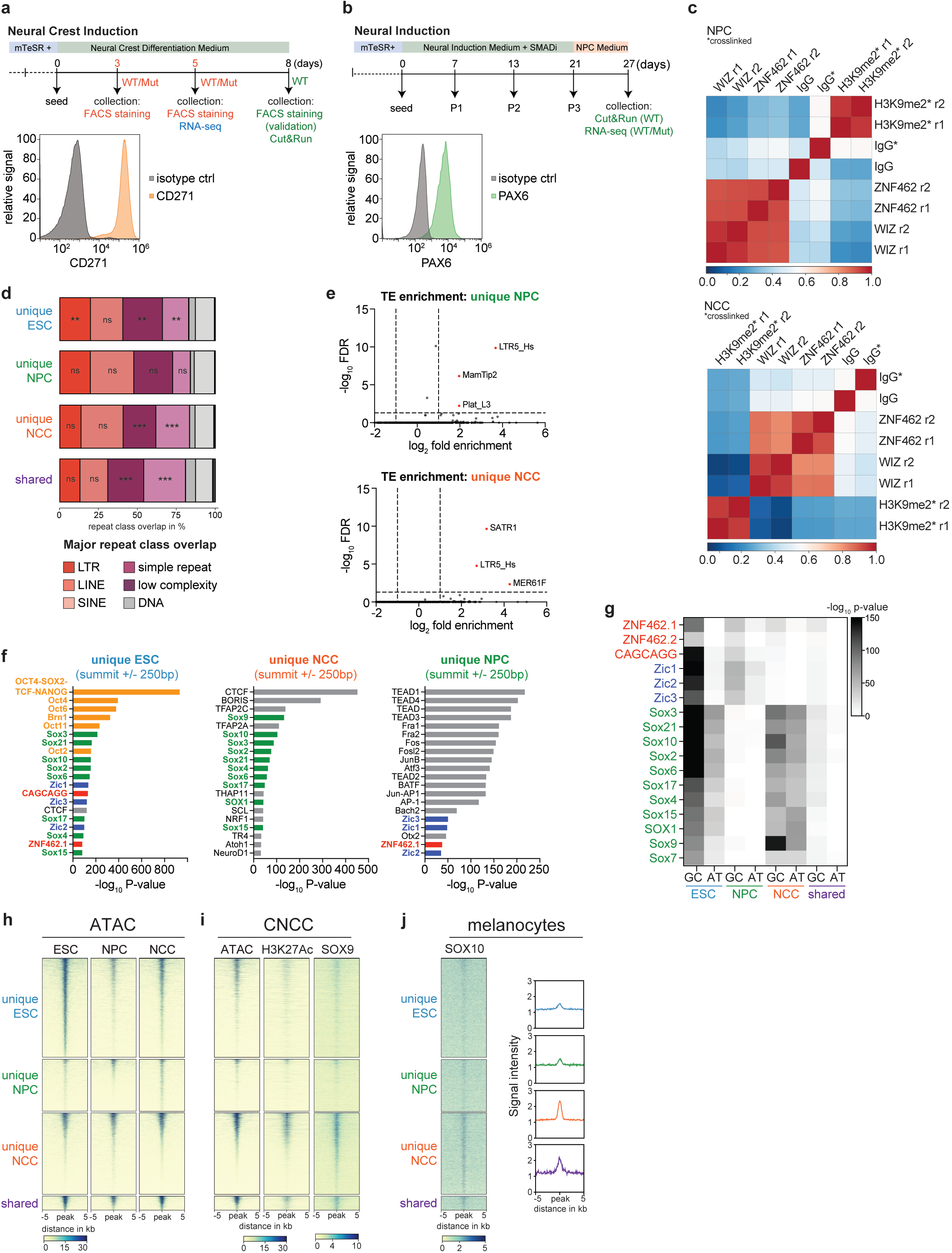
ZNF462 targets G9A/GLP to cell-type specific TE-Enhancers. a, **b)** Schematics of neural crest cell (NCC, left) and neural progenitor cell (NPC, right) differentiation protocols, including analysis time points and representative flow-cytometry validation of CD271 and PAX6 expression, respectively. **c)** Pearson correlation matrices of ZNF462, WIZ, H3K9me2, and matched IgG controls CUT&RUN datasets in WT NPCs and NCCs. Crosslinked samples are indicated by an asterisk (*). **d)** Distribution of major repeat-element classes overlapping unique ESC, unique NPC, unique NCC, and shared ZNF462 target sites. Statistical significance was assessed by one-sided Fisher exact test. ns, not significant; ** p < 0.01, *** p < 0.001. **e)** Volcano plots showing repeat-element subfamily enrichment at unique NPC and unique NCC ZNF462 target sites. The x-axis indicates log_2_ fold enrichment and the y-axis −log_10_ FDR. LTR5_Hs is significantly enriched in both unique NPC and unique NCC ZNF462 target sets. **f)** Bar plots showing the top 20 HOMER known TF motifs enriched at unique ESC, unique NPC, and unique NCC ZNF462 target sites. Motifs are ranked by HOMER enrichment P value and colored by motif family: POU (orange), SOX (green), ZIC (blue), ZNF462 (red), and other (gray). **g)** Heatmap comparing enrichment of selected motifs at unique ESC, unique NPC, unique NCC, and shared ZNF462 target sites using GC-matched genomic (GC) and accessibility-matched (AT) backgrounds. Values represent −log_10_ P value. **h)** Heatmaps of ATAC-seq signal at unique ESC, unique NPC, unique NCC, and shared ZNF462 target sites in hESCs, NPCs, and NCCs. **i)** Heatmaps of ATAC-seq, H3K27ac, and SOX9 signal at the same ZNF462 target classes in cranial neural crest cells^29^. **J)** Heatmap and metaplots of SOX10 signal at the same ZNF462 target classes in transformed melanocytes^30^.

**Ext. Data Fig. 5.**
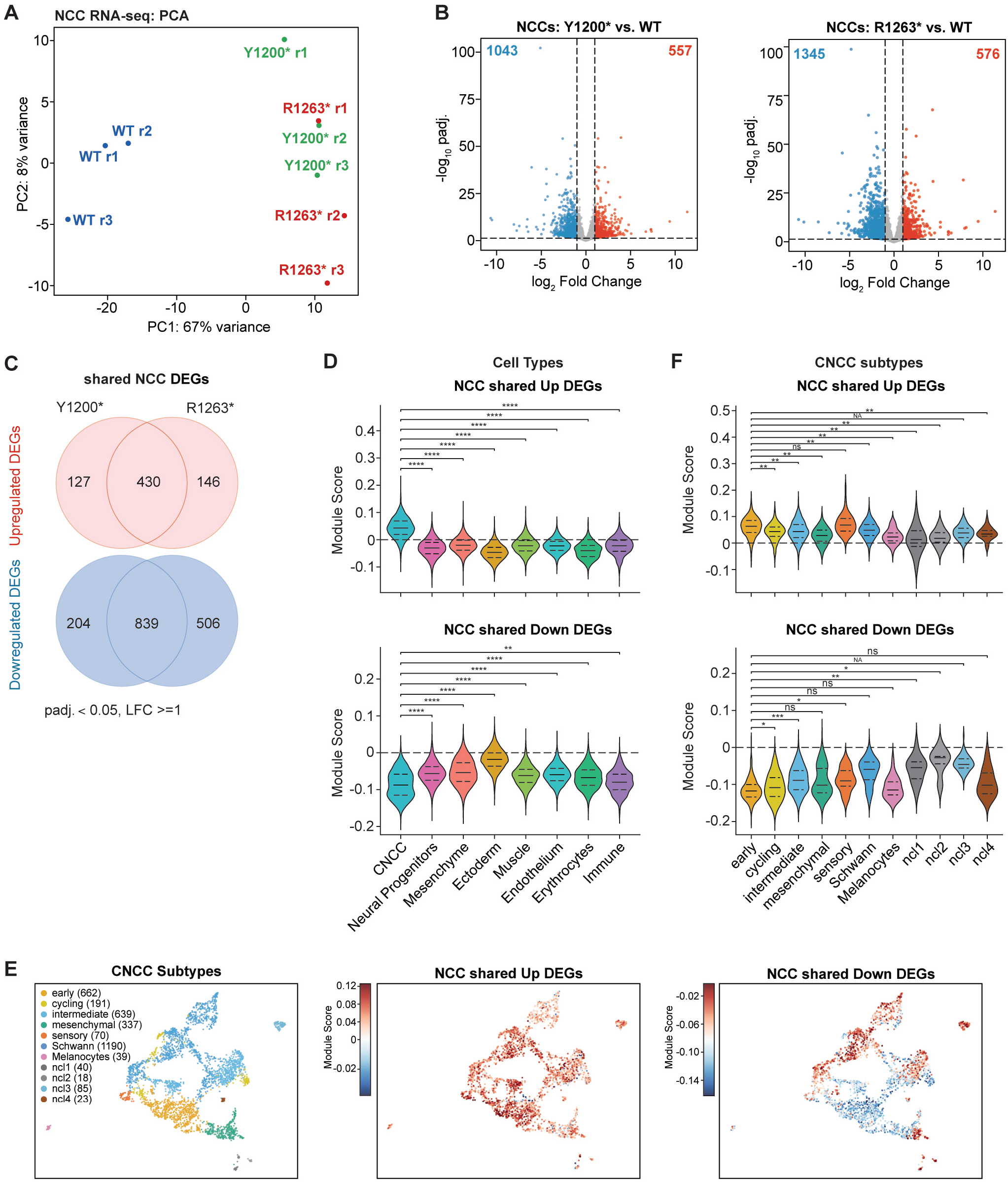
ZNF462 restricts timing of neural crest cell lineage specification. **a)** Principal component analysis (PCA) of RNA-seq expression in WT, Y1200*, and R1263* cells at day 5 of NCC differentiation. Each point represents an independent biological replicate. **b)** Volcano plots showing differential gene expression in NCCs derived from Y1200* and R1263* hESCs relative to WT controls. Up- and downregulated DEGs (FDR < 0.05, |log_2_ fold change| ≥ 1) are highlighted in red and blue, respectively. **c)** Venn diagrams showing the overlap of significantly upregulated and downregulated DEGs between Y1200* and R1263* NCCs relative to WT (FDR < 0.05, |log_2_ fold change| ≥ 1). **d)** Violin plots showing the distributions of module scores for the NCC shared-up (top) and NCC shared-down (bottom) gene signatures across eight cell types in the human craniofacial single-cell atlas (95,892 cells). Inner lines indicate quartiles. Statistical inference was performed by calculating the median module score for each cell population within each donor, followed by paired two-sided Wilcoxon signed-rank tests between the indicated reference population and comparison populations. P values were adjusted separately for Up and Down signatures using the Benjamini–Hochberg method. ns, non-significant, NA, not applicable, padj. ≥ 0.05; *, padj. < 0.05; **, padj. < 0.01; ***, padj. < 0.001; ****, padj. < 0.0001. **e)** UMAP projection of 3,294 cranial neural crest cells (CNCCs) from human craniofacial single-cell atlas^31^, colored by CNCC subtype (left) or by module scores for the NCC shared-up (middle) and NCC shared-down (right) gene signatures derived from ZNF462 mutant versus WT NCC bulk RNA-seq. CNCC subtypes comprise early (n = 662), cycling (n = 191), intermediate (n = 639), mesenchymal (n = 337), sensory (n = 70), Schwann (n = 1,190), melanocytes (n = 39), and neural crest-like populations ncl1 (n = 40), ncl2 (n = 18), ncl3 (n = 85), and ncl4 (n = 23). **f)** Violin plots showing the distributions of module scores for the NCC shared-up (top) and NCC shared-down (bottom) gene signatures across the 11 CNCC subtypes shown in E. Inner lines indicate quartiles. Statistical inference was performed by calculating the median module score for each cell population within each donor, followed by paired two-sided Wilcoxon signed-rank tests between the indicated reference population and comparison populations. P values were adjusted separately for Up and Down signatures using the Benjamini–Hochberg method. ns, non-significant, NA, not applicable, padj. ≥ 0.05; *, padj. < 0.05; **, padj. < 0.01; ***, padj. < 0.001; ****, padj. < 0.0001.

**Ext. Data Fig. 6.**
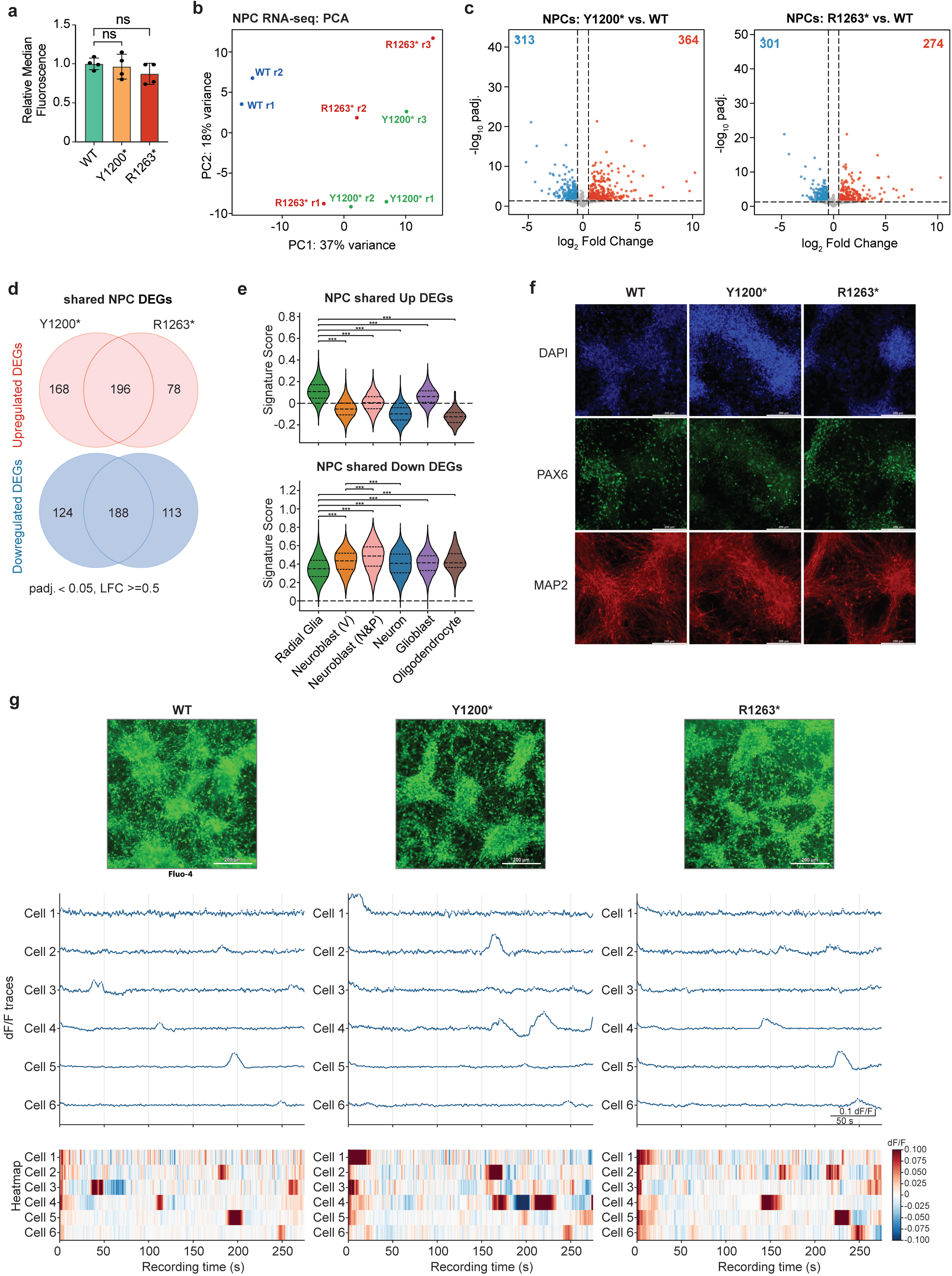
ZNF462 is required for proper timing of neuronal cell lineage specification. **a)** Quantification of PAX6 expression in WT, Y1200*, and R1263* cells following NPC differentiation, shown as relative median fluorescence. ns, not significant. **b)** Principal component analysis (PCA) of RNA-seq expression in WT, Y1200*, and R1263* cells at day 27 of NPC differentiation. Each point represents an independent biological replicate. **c)** Volcano plots showing differential gene expression in Y1200* and R1263* mutant NPCs relative to WT NPCs. Up- and downregulated DEGs (FDR < 0.05 and |log_2_ fold change| ≥ 0.5) are highlighted in red and blue, respectively. **d)** Venn diagrams showing the overlap of significantly upregulated and downregulated DEGs between Y1200* and R1263* NPCs relative to WT (FDR < 0.05 and |log_2_ fold change| ≥ 0.5). **e)** Violin plots showing the distributions of module scores for the NPC shared-up (top) and NPC shared-down (bottom) DEG signatures across six cell types in the first-trimester human telencephalon single-cell atlas (49,997 cells)^33^. Inner lines indicate quartiles. Statistical inference was performed by calculating the median module score for each cell population within each donor, followed by paired two-sided Wilcoxon signed-rank tests between the indicated reference population and comparison populations. P values were adjusted separately for Up and Down signatures using the Benjamini–Hochberg method. ns, non-significant, NA, not applicable, padj. ≥ 0.05; *, padj. < 0.05; **, padj. < 0.01; ***, padj. < 0.001; ****, padj. < 0.0001. **f)** Representative immunofluorescence images of cortical neurons differentiated from WT, Y1200*, and R1263* cells stained for PAX6 and MAP2. Nuclei were counterstained with DAPI. **g)** Representative Fluo-4 calcium-imaging fields, individual-cell ΔF/F traces, and corresponding activity heatmaps from cortical neurons differentiated from WT, Y1200*, and R1263* cells.

## Methods

### hESC culture conditions

H9 human embryonic stem cells (hESCs, WiCell WA09) were maintained on tissue culture plates coated with Corning® Matrigel® hESC-Qualified Matrix (Cat. No. 354277) in mTeSR Plus medium (STEMCELL Technologies, Cat. No. 100-0276). For recovery after thawing, 10 µM Y-27632 (STEMCELL Technologies, Cat. No. 72302) was supplemented in the culture medium for the first 24 hours. Cultures were maintained at 5% CO2 and 37°C with medium changes every 48 hours and passaged at 80% confluency, typically every 3–4 days. For routine maintenance, cells were passaged using ReLeSR (STEMCELL Technologies, Cat. No. 100-0483) at a 1:10 split ratio. For downstream assays requiring single-cell suspensions, hESCs were dissociated using ACCUMAX™ (STEMCELL Technologies, Cat. No. 07921) for 5 min incubation at 37°C.

### Generation of *ZNF462* mutant hESCs

Endogenous *ZNF462* mutant H9 hESC lines were generated by the IMBA Stem Cell Core Facility (ISCCF) using CRISPR/Cas9 ribonucleoprotein (RNP)-based genome editing. Two independent heterozygous nonsense mutant clones were generated, each harboring a single-nucleotide substitution in exon 3 of *ZNF462*. The Y1200* clone contains a heterozygous T-to-A substitution that introduces a premature stop codon near previously reported WKS patient mutations. The R1263* clone contains a heterozygous C-to-T substitution corresponding to the WKS-associated nonsense variant c.3787C>T, p.(Arg1263*) in ZNF462 (NM_021224.4), which introduces a premature stop codon. For genome editing, the Y1200* clone was generated using sgRNA462_1 (5′-CTGTGATTATGGGAACCGGA-3′), and the R1263* clone was generated using sgRNA462_3 (5′-AACTCCGAGCACTCAAATGT-3′). Edited clones were genotyped by PCR amplification of genomic DNA surrounding the targeted region using primers YR598 (5′-ACAGGCTCCTCCCACAGCT-3′) and YR599 (5′-CAGCTTCTATCAAGCCATCTAG-3′), which amplify a 488 bp product. PCR amplicons were gel-extracted and submitted to GENEWIZ (Azenta Life Sciences) for Sanger sequencing using the reverse sequencing primer YR645 (5′-GACGTGACTGTGGCTTTCAG-3′) to confirm the intended heterozygous single-nucleotide substitutions.

### Nuclear Protein Extraction

Single cells were collected, and nuclei were extracted by incubation in swelling buffer containing 10 mM Tris-HCl pH 7.5, 2 mM MgCl₂, 3 mM CaCl₂, and freshly added 1X Halt Protease Inhibitor Cocktail (Thermo Fisher Scientific, Cat. No. 78430) for 20 min with rotation at 4°C. Cells were pelleted by centrifugation at 500 × g for 5 min at 4°C, and the pellet was resuspended in cold GRO lysis buffer containing 10 mM Tris-HCl pH 7.5, 2 mM MgCl₂, 3 mM CaCl₂, 0.5% IGEPAL, 10% glycerol, and freshly added 1X Halt Protease Inhibitor Cocktail for 10 min with rotation at 4°C. Extracted nuclei were pelleted by centrifugation at 700 × g for 5 min at 4°C and lysed in RIPA buffer containing 150 mM NaCl, 5 mM EDTA pH 8.0, 50 mM Tris-HCl pH 8.0, 1% NP-40/IGEPAL CA-630, 0.5% sodium deoxycholate, and 0.1% SDS. RIPA buffer was freshly supplemented with 1 mM DTT, 1 mM PMSF, and 1X Halt Protease Inhibitor Cocktail. Benzonase nuclease was added to the lysate at a 1:100 dilution (Benzonase® Nuclease, Cat. No. 70746-4), followed by incubation on ice for 10 min. Lysates were further disrupted by passage through a 21-gauge needle and syringe. Lysates were then sonicated using a Bioruptor Pico (Diagenode, 20-60 kHz, 25-210 W) for 4 cycles of 15 sec ON and 30 sec OFF at 4°C. Lysates were centrifuged at 20,000 × g for 10 min at 4°C, and the supernatant containing the nuclear protein extract was collected for Western blot analysis.

### Western Blot analysis

Nuclear protein lysate (30 µg) was resolved on NuPAGE 3–8% Tris-Acetate mini protein gels (Invitrogen, EA0378BOX) using 1X NuPAGE Tris-Acetate SDS Running Buffer (Invitrogen, LA0041). Proteins were transferred onto Immobilon-P PVDF membranes (Millipore Sigma, Cat. No. IPVH00010). Membranes were blocked for 45 min at room temperature in 5% non-fat dry milk prepared in Tris-buffered saline with 0.1% Tween-20 (TBS-T). Membranes were then incubated overnight at 4°C with primary antibodies diluted in 5% non-fat dry milk in TBS-T. The primary antibodies used for Western blotting were rabbit anti-ZNF462 (1:1,000; Invitrogen, Cat. No. PA5-54585) and rabbit anti-TBP (1:1,000; Cell Signaling Technology, Cat. No. 8515S). Signal was detected using goat anti-rabbit IgG (H+L) DyLight 800 4X PEG secondary antibody diluted in 5% non-fat dry milk in TBS-T (1:10,000, Invitrogen, Cat. No. SA5-35571) for 45 min at room temperature. Membranes were imaged using Odyssey CLx Near-Infrared Imaging System (LICOR), and Western Blot signal was quantified using Image Studio Lite (LICOR) software.

### Flow cytometry analysis

Flow cytometry staining was performed on dissociated single-cell suspensions of hESCs, neural crest cells (NCCs), and neural progenitor cells (NPCs). For hESC pluripotency marker analysis, staining was performed using the Human/Mouse Pluripotent Stem Cell Multi-Color Flow Cytometry Kit (R&D Systems, Cat. No. FMC001) according to the manufacturer’s instructions. Briefly, hESCs were dissociated into single cells, washed twice with DPBS, and fixed using Fixation/Permeabilization Buffer for 30 min at 4°C. Fixed cells were resuspended in Permeabilization/Wash Buffer, and 10 µL of target antibody or corresponding isotype control was added to each sample. Samples were incubated for 45 min at room temperature protected from light, washed twice with Permeabilization/Wash Buffer, and resuspended in DPBS for flow cytometry analysis. For NCC and NPC marker analysis, dissociated single cells were first stained for the cell surface marker CD271. Cells were resuspended in Cell Staining Buffer (BioLegend, Cat. No. 420201) and incubated with anti-human CD271 Alexa Fluor 700 antibody (1:50, BioLegend, Cat. No. 345118) or corresponding isotype control for 30 min at room temperature protected from light. Following surface staining, cells were fixed in True-Nuclear Fix Buffer (BioLegend, Cat. No. 424401) for 1 hour at room temperature. Fixed cells were permeabilized and stained with anti-human PAX6-PE antibody (1:50, R&D Systems, Cat. No. IC8150P-100) or corresponding isotype control diluted in True-Nuclear Permeabilization Buffer (BioLegend, Cat. No. 424401) for 30 min at room temperature protected from light. Samples were washed and resuspended in True-Nuclear Permeabilization Buffer for flow cytometry analysis. Flow cytometry data were collected on an Attune NxT Acoustic Focusing Cytometer using Attune Cytometric Software (v7.1.3400.1). Final data analysis was performed using FlowJo (v10.10.1).

### Neural progenitor cell (NPC) differentiation

Neural progenitor cell differentiation was performed using the STEMdiff SMADi Neural Induction Kit (STEMCELL Technologies, Cat. No. 08581) according to the manufacturer’s monolayer differentiation protocol. Briefly, hESCs were dissociated into single cells and seeded onto Matrigel-coated 12-well plates at 1.4 × 10⁶ cells/well in Neural Induction Medium (NIM) supplemented with 10 µM Y-27632 for the first 24 hours. Medium was changed daily with fresh NIM until day 7. On day 7, cells were dissociated into single cells using ACCUMAX and replated onto Matrigel-coated 12-well plates at 1.4 million cells/well in NIM supplemented with 10 µM Y-27632 for the first 24 hours after passaging. Cultures were maintained with daily NIM changes for an additional 6 days. On day 13, cells were passaged a second time using the same dissociation and replating strategy. After an additional 8 days of culture with daily NIM changes, cells were passaged for a third time on day 21 and seeded onto Matrigel-coated 12-well plates at 1.4 million cells/well in Neural Progenitor Medium (NPM) supplemented with 10 µM Y-27632 for the first 24 hours. Cultures were maintained in NPM with daily medium changes for 6 days, and cells were collected on day 27 for downstream assays. Differentiation efficiency was validated in WT cells by flow cytometry analysis of PAX6 expression on day 27.

### Neural crest cell (NCC) differentiation

Neural crest cell differentiation was performed using the STEMdiff Neural Crest Differentiation Kit (STEMCELL Technologies, Cat. No. 08610) according to the manufacturer’s protocol. Briefly, hESCs were dissociated into single cells and seeded onto Matrigel-coated 6-well plates at 1 × 10⁶ cells/well in Neural Crest Differentiation Medium (NCDM) supplemented with 10 µM Y-27632 for the first 24 hours. Medium was changed daily with fresh NCDM until cells were collected for downstream assays. Differentiation efficiency was validated in WT cells by flow cytometry analysis of CD271 expression on day 8.

### Cortical neuron differentiation and immunostaining

hESCs were differentiated into cortical neurons following a previously published protocol^37^. Cells were cultured in CDM4 for 4 days after plating, followed by BrainPhys (STEMCELL Technologies, Cat. No. 05791) until day 11. Following calcium imaging acquisition, the same cultures were fixed in 4% paraformaldehyde (PFA) in PBS for 15 min at room temperature and washed three times with PBS. Cells were permeabilized with 0.1% Triton X-100 in PBS for 10 min and blocked in 4% BSA in PBS for 1 h at room temperature. Cultures were then incubated overnight at 4°C with primary antibodies (we need antibodies and the concentrations) diluted in blocking buffer. After three washes in PBS, cells were incubated for 1 h at room temperature, protected from light, with the appropriate Alexa Fluor–conjugated secondary antibodies (we need antibodies and the concentrations) diluted in blocking buffer. Nuclei were counterstained with DAPI, and cells were washed three times in PBS before imaging. Images were acquired on a Leica Thunder Imaging System.

### Calcium imaging of cortical neurons

For calcium imaging, differentiated neurons were incubated with Fluo-4 for 20 mins (Thermo Fisher Scientific, Cat. No. F14201). Upon PBS wash, cells were transferred to a recording chamber in BrainPhys Imaging Optimized Medium (STEMCELL Technologies, Cat. No. 05796). Calcium activity was recorded with a 20x objective, Leica Thunder Imaging System. Time-lapse images were acquired at 1frame every 200 ms for cortical neurons. Calcium imaging recordings (raw TIFF format) were analyzed in MATLAB (MathWorks) with the CaImAn toolbox^38^, applying constrained non-negative matrix factorization (CNMF) to extract regions of interest, identify fluorescent transients, and infer spike activity. Calcium traces are presented as relative scaled height as a function of time. Data are presented as mean ± SD. Statistical significance was determined using two-tailed t-tests or ANOVA with post hoc correction (\**p <* 0.05, \*\**p <* 0.01, \*\*\**p <* 0.001, \*\*\*\**p <* 0.0001).

### RNA-seq

Total RNA was extracted from cells using the Monarch Total RNA Miniprep Kit (New England Biolabs, Cat. No. T2010S) with on-column DNase digestion. For hESC RNA-seq, total RNA samples were submitted to GENEWIZ/Azenta Life Sciences for poly(A)-selected library preparation and sequencing on an Illumina NovaSeq platform with paired-end 150 bp reads. For NPC and NCC RNA-seq, total RNA samples were submitted to the USC Molecular Genomics Core for library preparation using the NEBNext Ultra II Directional RNA Library Prep Kit with poly(A) mRNA isolation (New England Biolabs, Cat. No. E7760S), followed by sequencing on a NextSeq 2000 platform with single-end 100 bp reads.

### CUT&RUN-seq

CUT&RUN was performed using the CUTANA™ ChIC/CUT&RUN Kit (EpiCypher, Cat. No. 14-1048) according to the manufacturer’s protocol, with minor modifications as described below. Cell permeabilization was performed using the protocol-recommended 0.01% digitonin concentration. For H3K9me2 CUT&RUN-seq, cells were lightly crosslinked with 0.1% formaldehyde for 1 min at room temperature, followed by quenching with glycine at a final concentration of 125 mM. For lightly crosslinked samples, the cell permeabilization buffer and antibody buffer provided in the kit were supplemented with 1% Triton X-100 and 0.05% SDS. Following cell collection, or following crosslinking where applicable, nuclei were extracted using nuclear extraction buffer based on the CUTANA™ ChIC/CUT&RUN Kit formulation, freshly supplemented with 0.5 mM spermidine and 1× Halt Protease Inhibitor Cocktail (Thermo Fisher Scientific, Cat. No. 78430). For CUT&RUN experiments in WT and mutant hESCs, *Drosophila* nuclei spike-in control (Active Motif, Cat. No. 53183) was added to the extracted nuclei according to the manufacturer’s instructions. Unless otherwise indicated, antibodies were added at 0.5 µg per reaction in 100 µL antibody buffer and incubated overnight. For the WT hESC ZNF462 CNR202405 CUT&RUN-seq dataset, ZNF462 antibody was used at either 0.2 µg or 0.5 µg per reaction. The following antibodies were used: H3K9me2 (Abcam, ab1220), ZNF462 (Invitrogen, PA5-54585), WIZ (Novus Biologicals, NBP1-80586), and G9A/EHMT2 (Bethyl Laboratories, A300-933A). Targeted chromatin digestion was performed at 4°C for 1 h for ZNF462, WIZ, and G9A/EHMT2, and for 2 h for H3K9me2. CUT&RUN-enriched DNA was used for library preparation with the CUTANA™ CUT&RUN Library Prep Kit (EpiCypher, Cat. No. 14-1001). Libraries were submitted to GENEWIZ (Azenta Life Sciences) for paired-end 150 bp sequencing on an Illumina platform.

### ATAC-seq

ATAC-seq was performed using the Active Motif ATAC-Seq Kit (Active Motif, Cat. No. 53150) according to the manufacturer’s protocol. Tn5 transposition was performed for 30 min for hESC samples and 15 min for NPC samples. Libraries were submitted to the Technology Center for Genomics & Bioinformatics at UCLA for paired-end 50 bp sequencing on an Illumina NovaSeq X Plus platform.

### RNA-seq data analysis

RNA-seq reads were quantified using Salmon against human GENCODE/GRCh38 v39 transcriptome indices^39^. Salmon quantifications were imported into R using tximeta and summarized to gene-level counts^40^. Differential expression analysis was performed using DESeq2 with a design formula of ∼ condition, with WT set as the reference condition^41^. Pairwise comparisons were performed for each ZNF462 mutant relative to WT. Genes with missing values were removed from downstream differential expression result tables. For downstream visualization and gene-list generation, genes with baseMean ≤10 were excluded. Differentially expressed genes were defined using an FDR cutoff of 0.05 and the indicated log_2_ fold-change thresholds. For quality control and visualization, variance-stabilizing transformed (VST) counts generated using DESeq2 were used for principal component analysis and heatmap visualization. PCA was performed using the top 500 most variable genes. For gene-expression heatmaps, VST-normalized expression values were row-scaled by gene to generate Z-scores across samples. For genome browser visualization, RNA-seq reads were independently aligned to the human GRCh38 reference genome using STAR with gene annotations from GENCODE v39^42^, and CPM-normalized bigWig tracks were generated from aligned BAM files using the bamCoverage function from deepTools with 10-bp bins and 30-bp smoothing^43^.

### CUT&RUN data processing and analysis

CUT&RUN sequencing reads were adapter-trimmed using Trim Galore and aligned with Bowtie2 (v2.5.2) to a hybrid genome consisting of human GRCh38, E. coli, and Drosophila melanogaster (dm6) sequences^44^. Aligned reads were filtered to retain high-quality mapped reads, remove PCR duplicates, and exclude CUT&RUN problematic regions^45^. ZNF462 peaks were called with MACS2 using matched IgG controls^46^, and reproducible ZNF462-bound regions were identified by Irreproducible Discovery Rate (IDR) analysis (https://doi.org/10.1214/11-AOAS466). The resulting conservative IDR peak set from the CNR202405 dataset was used as the high-confidence ZNF462 reference peak set for downstream analyses.

Normalized CUT&RUN-seq signal tracks were generated from filtered BAM files using deepTools and used for genome browser visualization, heatmaps, metaplots, and quantitative signal analyses. WT CUT&RUN-seq datasets were normalized using reads-per-genomic-content (RPGC) normalization, whereas WT and mutant CNR202501 datasets were normalized using Drosophila spike-in scaling. Sample quality and replicate concordance were assessed using genome-wide Pearson correlation matrices generated from normalized signal tracks. T1 lamina-associated domain (LAD) coordinates, representing LADs with the highest Lamin B1 enrichment, were obtained from Shah et al.^47^. Quantitative CUT&RUN-seq signal over ZNF462-bound regions and LADs was measured using normalized ZNF462 and H3K9me2 bigWig tracks. High-confidence ZNF462 peak-centered regions and T1 LAD-centered regions were defined as mutually exclusive region sets, and signal distributions were quantified using bigWigAverageOverBed and visualized as violin plots.

### Transcription factor occupancy and motif analyses

Previously published hESC SOX2 CUT&RUN (SRP188770)^48^ and hESC OCT4 ChIP-seq (SRP059231)^49^ datasets were reprocessed from raw sequencing reads. Reads were aligned to the human GRCh38 reference genome, peaks were called using the corresponding matched control samples, and reproducible SOX2 and OCT4 peak sets were defined using IDR analysis. Previously published ZIC2 ChIP-seq data from primed human pluripotent stem cells (SRP516114) ^50^ were also reprocessed from raw sequencing reads. Adapter sequences were removed using Cutadapt, reads were aligned to the human GRCh38 reference genome using Bowtie2 (v2.5.2), and alignments with mapping quality <30 were removed. PCR duplicates were marked and excluded using Picard MarkDuplicates and reads overlapping hg38 blacklist v2 regions (Boyle Lab blacklist collection)^51^ or noncanonical chromosomes were removed. RPGC-normalized ZIC2 signal tracks were generated from filtered BAM files using deepTools bamCoverage, and replicate-specific peaks were called using MACS2 with the corresponding matched input samples as controls. For downstream co-occupancy analyses, the published ZIC2 peak set provided with the original study was used. ZNF462, OCT4, SOX2, and ZIC2 peak sets were compared using Intervene^52^ with BEDTools-based interval overlap^53^. Overlap combinations were consolidated into four mutually exclusive region classes: ZNF462-specific regions; ZNF462-shared regions overlapping at least one of OCT4, SOX2, or ZIC2; OCT4–SOX2 regions lacking ZNF462, with or without ZIC2 co-occupancy; and ZIC2-containing regions lacking ZNF462 that were not assigned to the OCT4–SOX2 class. Normalized ZNF462, OCT4, SOX2, and ZIC2 signal across these region classes was visualized using heatmaps and metaplots generated with deepTools. For early NCC analyses, previously published day 5 NCC SOX2 CUT&RUN data from Hovland et al. (2022) (SRP299576)^54^ were reprocessed from raw sequencing reads using the same general CUT&RUN processing workflow and used for comparative occupancy analyses at ZNF462-bound regions. ZNF462 overlap with candidate cis-regulatory elements was assessed using SCREEN Registry candidate cis-regulatory element (cCRE) annotations^25^. ZNF462 peaks were intersected with SCREEN cCREs using BEDTools and assigned to representative cCRE classes. H9-core SCREEN cCRE annotations were used to define pluripotency-context enhancer- and promoter-like regulatory elements and to compare ZNF462-bound elements with matched unbound regulatory elements. H3K27ac, H3K4me1, and H3K4me3 datasets from H9 hESCs were obtained from the 0-h time point of SRP234480^55^. Raw sequencing reads were processed and aligned to the human GRCh38 reference genome, and RPGC-normalized signal tracks were generated for downstream analyses. Gene-linking analyses were performed by intersecting ZNF462-bound enhancer cCREs with selected SCREEN v4 computational gene-link annotations from developmental and pluripotency contexts. High-confidence links were defined as those supported by at least two SCREEN computational methods or by EPIraction and were used for integration with differential expression and accessibility analyses. Gene ontology enrichment of linked genes was performed using AmiGO 2 with the GO Biological Process Complete annotation set, with all human genes as the reference background and FDR correction for multiple testing^56^.

Transcription factor motif enrichment at ZNF462-bound regions was evaluated using HOMER^57^ in 250-bp and 500-bp summit-centered windows. Analyses used both GC-matched genomic backgrounds and GC-matched accessibility-controlled backgrounds centered on lineage-matched WT ATAC-seq summits. hESC, NPC, and NCC peak sets were compared with hESC WT, NPC WT, and NCC day 5 ATAC-seq (SRP299102)^54^ backgrounds, respectively, whereas regions shared among cell types were compared with a pooled hESC/NPC/NCC ATAC-seq background. HOMER analyses used a vertebrate motif collection supplemented with the ZNF462 motifs UN0616.1 and UN0616.2, and AME analyses used the JASPAR 2024 vertebrate CORE collection^58^ supplemented with the same ZNF462 motifs.

To assess the relationship between motif content and chromatin accessibility following ZNF462 loss, the 30 highest-ranking known motifs were selected independently from HOMER analyses of hESC ZNF462 peaks using 500-bp windows. ZNF462-bound regions were scanned for these motifs using HOMER, and motif occurrence was represented as a binary variable. For each motif, motif-positive and motif-negative regions were compared with respect to baseline WT accessibility and average accessibility change following ZNF462 loss. Associations between motif occurrence and accessibility change were assessed by Spearman correlation and Mann–Whitney U tests, with rank-biserial correlation used as an effect-size measure and Benjamini–Hochberg correction applied across motifs. The relationship between motif-associated baseline accessibility and accessibility gain was assessed by Spearman rank correlation across the 30 motifs independently for the 500-bp window size motif sets.

### Transposable element enrichment analysis

Repeat enrichment was assessed using ZNF462 peak intervals and RepeatMasker annotations from the UCSC hg38 rmsk table, restricted to chromosomes 1–22 and X. Subfamily analyses included LTR, LINE, SINE, DNA, and Satellite repeats represented by ≥10 genomic elements. Peak–repeat intersections were determined using BEDTools and deduplicated such that each peak was counted at most once per subfamily. Descriptive enrichment was calculated as the fraction of ZNF462 peaks overlapping each subfamily relative to its nonredundant genomic coverage. Statistical enrichment was assessed using 100 randomized genomic region sets matched to ZNF462 peaks by chromosome and interval length using BEDTools shuffle. Observed and matched-background overlap frequencies were compared using one-sided Fisher’s exact tests with Benjamini–Hochberg correction within each peak set. Effect sizes were reported as matched-background fold enrichment and Fisher odds ratios. Heatmaps display log2-transformed matched-background fold enrichment for significant subfamilies, while scatterplots show log2 fold enrichment versus −log_10_ FDR.

Repeat-class distributions were additionally evaluated across 11 RepeatMasker categories: LTRs, LINEs, SINEs, simple repeats, ncRNAs, low-complexity sequences, DNA transposons, satellites, scRNAs, snRNAs, and unknown repeats. Class-level overlap and enrichment were assessed for hESC, NPC, NCC, combined NPC+NCC, and hESC-only, NPC-only, NCC-only, and shared ZNF462 peak sets. Peak overlap was expressed as the percentage of peaks overlapping each category, and enrichment was calculated relative to the mean overlap across 100 chromosome- and interval-length-matched randomized sets, with significance assessed using one-sided empirical P values. For stacked composition plots, category-specific overlaps were normalized to the total overlaps across the 11 displayed categories.

### Single-cell reference atlas analysis

Previously published human single-cell RNA-seq reference atlases were used to evaluate the developmental and cell-type distribution of transcriptional signatures associated with *ZNF462* haploinsufficiency. For neural lineage analyses, the processed human developmental brain atlas from Braun et al. (2023) was analyzed while retaining the original cell annotations and embedding supplied with the dataset^33^. ZNF462 mutant NPC signatures were defined from genes differentially expressed concordantly in the Y1200* and R1263* lines relative to WT. Shared-upregulated and shared-downregulated gene sets were scored independently in individual atlas cells using scanpy.tl.score_genes^59^ with expression-matched control genes. Signature scores were visualized on the reference embedding and summarized across annotated neural cell classes. To account for non-independence of cells from the same individual, median signature scores were additionally calculated by donor and cell class, and paired comparisons between radial glia and other neural populations were performed using two-sided Wilcoxon signed-rank tests with Benjamini–Hochberg correction.

For neural crest lineage analyses, processed single-cell RNA-seq data from Khouri-Farah et al. (2026) were analyzed using the published cell annotations and UMAP coordinates^31^. Concordant ZNF462 mutant NCC transcriptional signatures were scored in individual cells using scanpy.tl.score_genes, with shared-upregulated and shared-downregulated signatures evaluated independently. Signature scores were visualized on the reference UMAP embeddings and summarized across annotated cell types and CNCC subpopulations.

### ATAC-seq data analysis

Raw paired-end ATAC-seq reads from hESC and NPC samples were trimmed for Nextera adapter sequences using Cutadapt (v2.10; DOI: 10.14806/ej.17.1.200) and aligned to a hybrid human GRCh38/hg38–Drosophila melanogaster dm6 reference genome using Bowtie2 (v2.5.2), allowing fragment sizes up to 2 kb and retaining up to four valid multimapping alignments per read. Previously published day 5 NCC ATAC-seq data from Hovland et al. (SRP299102)^54^ and committed CNCC ATAC-seq data from Naqvi et al. (SRP379721)^60^ were reprocessed from raw reads and aligned to GRCh38. Alignments were sorted using SAMtools (v1.19.2), and unmapped, improperly paired, and PCR-duplicate reads were removed. Tn5-corrected tagAlign files were generated using BEDTools by shifting read starts +4 bp and −5 bp on the positive and negative strands, respectively^61^. Accessible regions were identified using MACS2 (P < 0.01; no-model; shift −75 bp; extension 150 bp), reproducible peaks were determined by IDR analysis, and hg38 blacklist v2 regions were excluded (PMID: 31249361). Differentially accessible regions (DARs) were identified from a nonredundant union of 500-bp summit-centered ATAC-seq peaks across WT and ZNF462-mutant hESCs. Paired-end fragments were quantified using BEDTools, and differential accessibility between WT and the Y1200* and R1263* lines was assessed independently using DESeq2, with log_2_ fold-change shrinkage using apeglm^62^. Concordance between mutant lines was assessed by Spearman correlation of log2 fold changes across all 162,954 ATAC-seq regions prior to significance filtering. DARs were defined by FDR < 0.05 and |log_2_FC| ≥ 0.5; regions showing concordant increases or decreases in both mutants were classified as Shared-Up or Shared-Down, respectively. For integrated analyses, the mean Y1200*/R1263* log2 fold-change was used as a summary measure of accessibility change. RPGC-normalized ATAC-seq bigWig tracks were generated from Tn5-shifted alignments using deepTools bamCoverage and used for visualization and quantitative accessibility analyses. Baseline accessibility was defined as mean WT ATAC-seq signal. The relationship between baseline accessibility and accessibility change among ZNF462-bound Shared-Up DARs was assessed using a two-sided Spearman rank correlation. To control for pre-existing chromatin accessibility in motif enrichment analyses, ZNF462-bound regions were compared with two matched backgrounds. Genomic control regions were matched one-to-one for GC content, whereas accessibility-controlled regions were selected from WT ATAC-seq peaks and jointly matched one-to-one, without replacement, for GC content and quantitative WT ATAC-seq signal. Regions overlapping ZNF462 targets were excluded from the matched backgrounds.

### Statistics and reproducibility

All experiments were independently repeated at least two times (unless otherwise stated) with similar results, and representative results are shown throughout all figures. No statistical method was used to predetermine sample size. No data were excluded from the analyses. The experiments were not randomized. The Investigators were not blinded to allocation during experiments and outcome assessment.

